# Rudimentary Structure-Activity-Relationship study of the MMV Pathogen Box compound MMV675968 (2,4-diaminoquinazoline) unveils novel inhibitors of Trypanosoma brucei brucei DHFR enzyme

**DOI:** 10.1101/2022.05.20.492762

**Authors:** Darline Dize, Rolland Tata, Rodrigue Keumoe, Rufin Marie Kouipou Toghueo, Mariscal Brice Tchatat, Cyrille Ngansop Njanpa, Vianey Claire Tchuenguia, Lauve Tchokouaha Yamthe, Patrick Valere Tsouh Fokou, Benoît Laleu, James Duffy, Ozlem Tastan Bishop, Fabrice Fekam Boyom

**Affiliations:** Antimicrobial and Biocontrol Agents Unit (AmBcAU), Laboratory for Phytobiochemistry and Medicinal Plants Studies, Department of Biochemistry, Faculty of Science, University of Yaoundé I, Yaoundé, Cameroon; Research Unit in Bioinformatics (RUBi), Department of Biochemistry and Microbiology, Rhodes University, Grahamstown, South Africa; Medicines for Malaria Venture, Route de Pré-Bois 20, Meyrin, Switzerland

**Author notes:** Corresponding author: Fabrice Fekam Boyom; Email address.

**Keywords:** *Trypanosoma brucei brucei*, MMV Pathogen Box, Antitrypanosomal, MMV675968 (2,4-diaminoquinazoline), *In silico*, Structure-activity-relationship, DHFR inhibitor, *time*-kill kinetic, DNA fragmentation

## Abstract

New drugs are urgently needed for the treatment of human African trypanosomiasis (HAT). In line with our quest for dihydrofolate reductase (DHFR) inhibitors in trypanosomes, a small library of analogs of the antitrypanosomal hit (**MMV675968**) available at MMV as solid materials was screened for antitrypanosomal activity. *In silico* confirmation of two potent antitrypanosomal analogs as inhibitors of DHFR was achieved, together with elucidation of other antitrypanosomal modes of action. Overall, two analogs of the diaminoquinazoline derivative, **MMV675968** reputed to inhibit the enzyme DHFR, displayed approximately 40-fold and 60-fold more potency and selectivity than the parent hit, respectively (**MMV1578445 (10):** IC_50_ = 0.045 µM, SI=1737; **MMV1578467 (7):** IC_50_ = 0.06 µM; SI=412). Analogs **7** and **10** were also strong binders of the DHFR enzyme, in all their accessible protonation states, and interacted with key DHFR ligand recognition residues Val32, Asp54, and Ile160. **MMV1578445 (10)** portrayed fast and irreversible trypanosomes growth arrest after 72 h and 4 h at IC_99_. Analogs **7** and **10** induced ferric iron reduction and DNA fragmentation or apoptosis induction, respectively. The two potent analogs endowed with suitable physicochemical properties are good candidates for further deciphering their potential as starting points for new drug development for HAT.

**Author summary:** African trypanosomiasis is a disease caused by parasites of the *Trypanosoma brucei* species which affect both humans and animals. We have adopted the fast-track drug repurposing approach to determine the antitrypanosomal activity of the open access MMV Pathogen Box compound library. The compound 2,4-diaminoquinazoline (MMV675998), known as a dihydrofolate reductase (DHFR) inhibitor was identified. Rudimentary structure-activity relationship investigation of this compound unveiled two highly active and selective derivatives MMV1578445 and MMV1578467 which were found to strongly inhibit the *Trypanosoma brucei* dihydrofolate reductase enzyme *in silico*. Besides, we demonstrated that analogues MMV1578445 and MMV1578467 could respectively reduce intrinsic ferric iron and induce apoptosis in trypanosomes. The two identified inhibitors of trypanosomes qualify as potential drug candidates that could be useful for future development as novel antitrypanosomal drugs.

## Introduction

African trypanosomiasis is a neglected tropical disease that mostly results from the bite of infected *tsetse* flies bearing flagellated kinetoplastid parasites of the genus *Trypanosoma* [1]. The disease is a huge threat to both humans (Human African Trypanosomiasis, HAT) and animals (Animal African Trypanosomiasis, AAT) with enormous social and economic impacts in endemic regions. HAT, also called sleeping sickness, is caused by two subspecies of *Trypanosoma brucei* sp.: *Trypanosoma brucei gambiense* in western and central Africa and *Trypanosoma brucei rhodesiense* found in eastern and southern Africa, which induce chronic and acute forms of the disease, respectively [2]. The former accounts for approximately 95% of the total number of reported cases, while the latter causes approximately 5%, with 65 million people being at risk of infection in the 36 endemic countries. Since 2009, the number of reported cases has dropped from 40 000 to 993 in 2018, leading to the launch of the programme toward its elimination by the World Health Organization (WHO) [3]. However, in 2019, unusual recrudescence was observed in foci of some endemic countries, such as Cameroon [4, 5], supporting the idea of pursuing effort for the global view of eradication rather than slackening disease surveillance. Animal African trypanosomiasis, also called nagana disease, caused by *Trypanosoma brucei brucei* remains the most important cattle disease in sub-Saharan Africa with severe economic consequences [6]. The disease considerably hampers livestock production by threatening approximately 50 million head of cattle with 3 million deaths per year [7]. Due to vaccine development failures, the ideal management tool used for decades is chemotherapy, which relies on the use of validated treatment options depending on parasite species and the disease stages. The inefficiency and toxicity of the current remedies combined with the development of resistance represent the main bottleneck of trypanosomiasis control [6, 8]. Hence, the development of novel and adequate treatment options is of high priority to contribute to the global aim of eradicating the disease.

Several strategies have been developed to identify new and requisite chemical entities for drug development toward parasitic infections. These include drug repositioning, where drugs already approved for a particular infection are reprofiled for another disease [9]. This method has been previously used in parasitic diseases. For instance, most of the leishmaniasis drug arsenal was repurposed from other indications. This is the case for marketed amphotericin B, originally used in fungal infections and later redirected for the treatment of leishmaniasis [9, 10]. Miltefosine is the most recent antileishmanial but initially developed for cutaneous metastases of breast cancer and solid tumors [11]. The antifungal poseconazole is a repurposed candidate that has reached the first clinical trials for Chagas disease [12]. Globally, drug repositioning is an interesting approach to rapidly provide new alternatives against neglected tropical diseases. Within this framework, US FDA very recently approved the use of Fexinidazole [13] that has been developed by DND*i* as the first oral treatment for sleeping sickness. This 2-substituted 5-nitroimidazole was primarily designed during the seventies in an attempt to development anti-infective agents [14].

Fortunately, sources of repositionable compounds have been facilitated and encouraged by the work of some organizations, such as the Medicines for Malaria Venture (MMV), through Open Source Drug Discovery. They have therefore assembled many collections of compounds directed against malaria and other diseases [15], among which the Pathogen Box (MMVPB) consisting of 400 diverse drug-like molecules active against neglected diseases [16]. The MMVPB has been successfully investigated, leading to the discovery of newly identified activity against various pathogens. For example, compounds MMV010576, MMV028694 and MMV676501 showed inhibitory activity at micromolar levels with dual efficacy toward *Giardia lamblia* and *Cryptosporidium parvum* [17]. On the other hand, the insecticide Tolfenpyrad (MMV688934) showed good activity against the helminth parasite and barber’s pole worm [18]. The anti-*Toxoplasma gondii* activity of the Pathogen Box library was investigated by [19], leading to the identification of eight selective hit compounds. **Duffy et al.** [20] also identified chemical starting points for malaria, human African trypanosomiasis, Chagas disease and leishmaniasis drug discovery from their *in vitro* antiprotozoal screening. Screening of the Pathogen Box against *Echinococcus multilocularis* identified the old drug buparvaquone (MMV689480) as having promising activity [21]. Similarly, we explored the Pathogen Box with the aim of identifying promising starting points for drug discovery against trypanosomiasis.

We herein present the activity profile of compounds emerging from the screening of the MMVPB against the bloodstream forms of *Trypanosoma brucei brucei*. A selected trypanocidal hit was further processed for a preliminary structure-activity relationship study using analogs available at MMV. The most potent analogs were analyzed for time-kill kinetics, reversibility of trypanocidal effect, effect on parasite plasma membrane integrity, measurement of reactive oxygen species, DNA fragmentation analysis and ferric iron reducing potency.

## Results

### Identification of pathogen box compounds as inhibitors of bloodstream forms of *Trypanosoma brucei brucei*

#### Preliminary screening of the MMV Pathogen Box

The primary screening of the 400 MMV PBox compounds at a fixed dose of 10 µM led to the identification of 70 compounds that inhibited the viability of trypanosomes by at least 90% (Fig 1). According to the MMVPB supporting information [16], 7 of the hits were reference compounds, including 5 anti-trypanosomatid drugs, *viz.* Two anti-HAT drugs (pentamidine and suramin), 2 anti-chagasic drugs (nifurtimox and benznidazole) and 1 antileishmanial drug (sitamiquine), thus validating the anti-trypanosomal assay performed, and 2 antimalarial drugs (mefloquine and primaquine) (Table 1). In addition, 25 MMVPB compounds reported to inhibit kinetoplastids (*Trypanosoma* and *Leishmania* spp.) were also identified (Table 2). The last group of inhibitors consisted of 38 compounds reported to target other diseases, including tuberculosis (16), malaria (14), schistosomiasis (03), toxoplasmosis (02), cryptosporidiosis (02), and filariasis (01) (**Fig 1B**).

**Fig 1.**
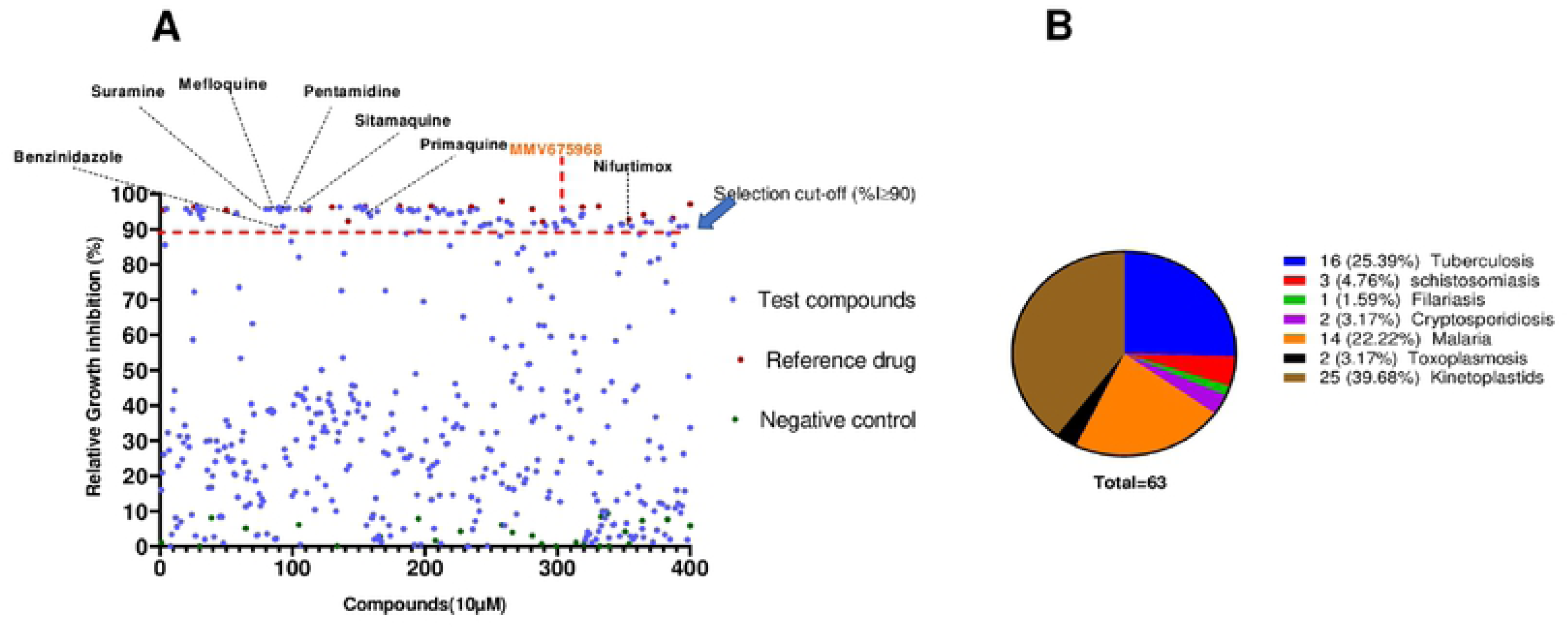
Antitrypanosomal activity cutoff of the 400 MMVPB compounds. (A) Compounds were screened at 10 µM using the resazurin assay, and inhibitory percentages were determined relative to the negative control culture. The 90% cutoff criterion (black dotted line) was used to select the most promising inhibitors for concentration–response assays. (B) Pie chart showing the distribution of selected inhibitors according to their reported disease targets.

**Table 1:**
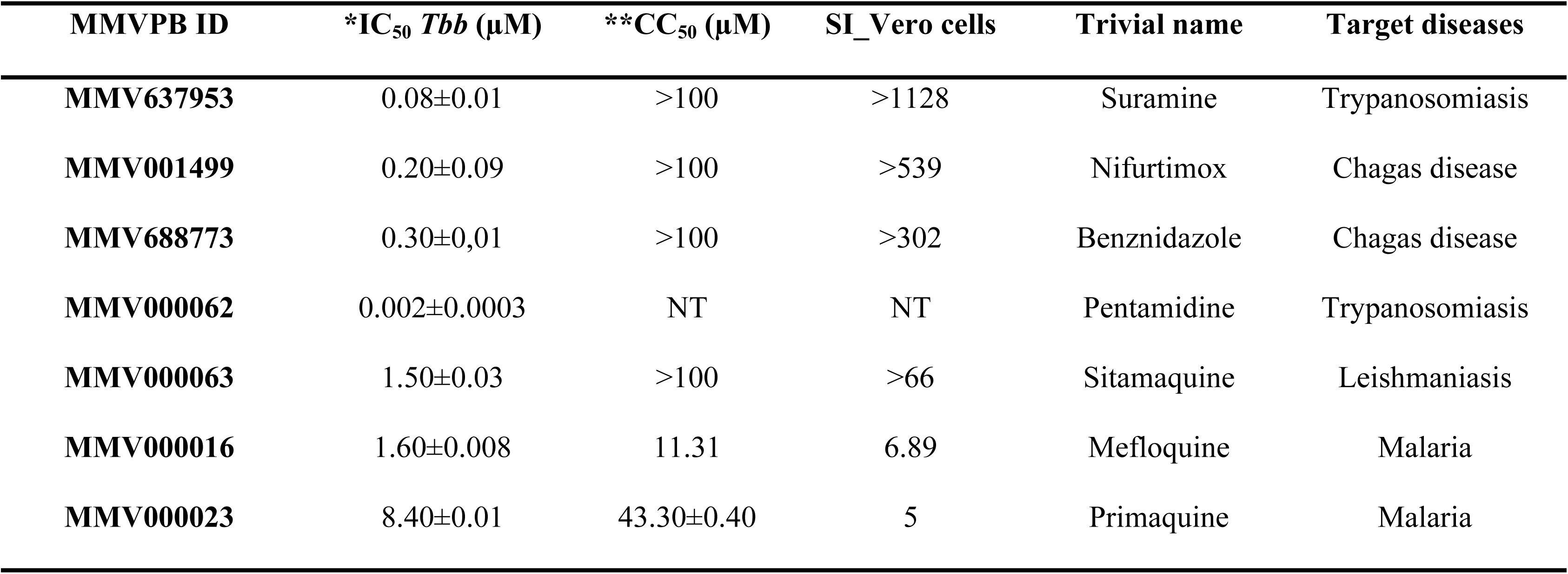
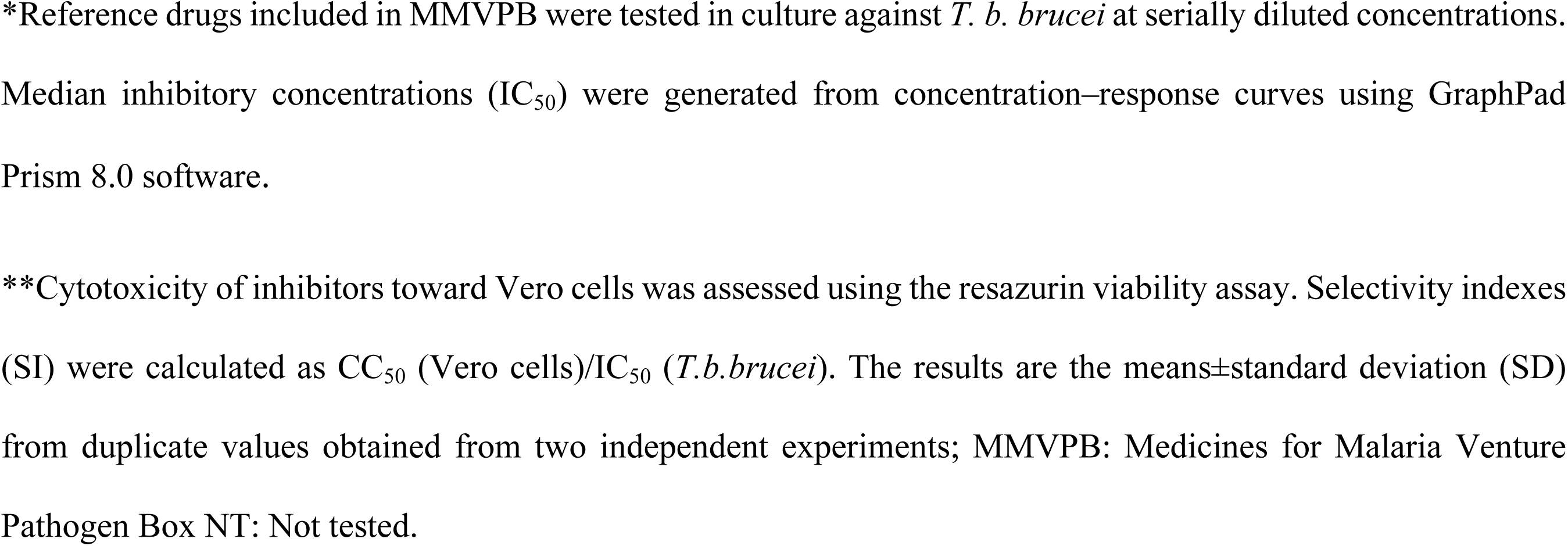
Antitrypanosomal activity of reference drugs included in the MMV Pathogen Box.

| MMVPB ID | *IC<sub>50</sub> *Tbb* (µM) | **CC<sub>50</sub> (µM) | SI_Vero cells | Trivial name | Target diseases |
| --- | --- | --- | --- | --- | --- |
| MMV637953 | 0.08±0.01 | >100 | >1128 | Suramine | Trypanosomiasis |
| MMV001499 | 0.20±0.09 | >100 | >539 | Nifurtimox | Chagas disease |
| MMV688773 | 0.30±0.01 | >100 | >302 | Benznidazole | Chagas disease |
| MMV000062 | 0.002±0.0003 | NT | NT | Pentamidine | Trypanosomiasis |
| MMV000063 | 1.50±0.03 | >100 | >66 | Sitamaquine | Leishmaniasis |
| MMV000016 | 1.60±0.008 | 11.31 | 6.89 | Mefloquine | Malaria |
| MMV000023 | 8.40±0.01 | 43.30±0.40 | 5 | Primaquine | Malaria |

\*Reference drugs included in MMVPB were tested in culture against *T. b. brucei* at serially diluted concentrations. Median inhibitory concentrations (IC<sub>50</sub>) were generated from concentration–response curves using GraphPad Prism 8.0 software.
\*\*Cytotoxicity of inhibitors toward Vero cells was assessed using the resazurin viability assay. Selectivity indexes (SI) were calculated as CC<sub>50</sub> (Vero cells)/IC<sub>50</sub> (*T.b.brucei*). The results are the means±standard deviation (SD) from duplicate values obtained from two independent experiments; MMVPB: Medicines for Malaria Venture Pathogen Box NT: Not tested.

**Table 2:** *In vitro* antitrypanosomal and cytotoxic activities of the 25 MMVPB compounds with known anti-kinetoplastid activity.

| Compound No | MMVPB ID | *IC<sub>50</sub> *T.b. brucei* (μM) | **CC<sub>50</sub><br>Vero | SI | †Chemical Classes | Trivial name | #Reported IC<sub>50</sub> (μM) |
| --- | --- | --- | --- | --- | --- | --- | --- |
| 1 | MMV688180 | 0.002±0.01 | >100 | >42826 | Benzenesulfonamide |  | <0.13 |
| 2 | MMV688796 | 0.03±0.01 | >100 | >3416 | 2,4 substituted furan |  | 0.1 |
| 3 | MMV676604 | 0.036±0.001 | 8.30±1.20 | 232 | 2-aminopyrimidine |  | 0.26 |
| 4 | MMV688797 | 0.06±0.02 | >100 | >1598 | 2-aryl oxazole |  | 0.13 |
| 5 | MMV652003 | 0.06±0.04 | >100 | >1557.87 | Benzamide |  | 0.15 |
| 6 | MMV688958 | 0.087±0.01 | >100 | >1154 | 2-aryl oxazole |  | 0.15 |
| 7 | MMV675998 | 0.34±0.03 | >100 | >296 | Benzenecarboximidamide |  | 0.25 |
| 8 | MMV688798 | 0.30±0.025 | NT | NT | Benzamide |  | 0.555 |
| 9 | MMV688795 | 0.35±0.02 | >100 | >282 | 2-aryl oxazole |  | 0.15 |
| 10 | MMV688793 | 0.36±0.04 | >100 | >255 | 2-pyridyl benzamides |  | 2.07 |
| 11 | MMV689028 | 0.40±0.02 | >100 | >248 | benzyl piperazine |  | 0.14 |
| 12 | MMV676600 | 0.65±0.007 | >100 | >154 | Benzamide |  | 1.02 |
| 13 | MMV188296 | 1.01±0.25 | >100 | >98 | 2-indolinecarboxamide |  | 0.49 |
| 14 | MMV688271 | 1.07±0.16 | >100 | >93 | Guanidine |  | 0.6 |
| 15 | MMV689029 | 1.30±0.60 | >100 | >76 | benzyl piperazine |  | 0.5 |
| 16 | MMV688371 | 1.60±0.10 | 7.5±0.1 | 5 | Benzamide |  | <0.13 |
| 17 | MMV689061 | 1.90±0.10 | >100 | >54 | Acetamide |  | 32.3 |
| 18 | MMV001561 | 1.90±0.05 | 41.5±1.0 | 22 | Propanamine | fluoxetine | 3.97 |
| 19 | MMV687706 | 1.90±0.006 | 41.0±6.5 | 21 | Piperazine |  | 0.99 |
| 20 | MMV659004 | 1.90±0.003 | >100 | >50 | Pyrimidine |  | 6.6 |
| 21 | MMV689060 | 2.40±0.06 | NT | NT | Piperazine |  | 31.6 |
| 22 | MMV690027 | 2.80±0.07 | >100 | >36 | hexahydrophthalazinones |  | 0.02 |
| 23 | MMV688467 | 3.10±0.50 | >100 | >32 | butyl sulfanilamide |  | 0.3 |
| 24 | MMV688514 | 3.20±0.30 | 17±2.5 | 5 | Benzenecarboximidamide |  | 4.06 |
| 25 | MMV688410 | 8.50±0.10 | NT | - | Acetamide | - | 3.8 |

\* MMVPB compounds were tested in culture against *T. b. brucei* at serially diluted concentrations. Median inhibitory concentrations (IC<sub>50</sub>) were generated from concentration– response curves using GraphPad Prism 8.0 software.
\*\*Cytotoxicity of inhibitors toward Vero cells was assessed using the resazurin viability assay. Selectivity indexes (SI) were calculated as CC<sub>50</sub> (Vero cells)/IC<sub>50</sub> (*T.b.brucei*).
The results are means±standard deviation (SD) from duplicate values obtained from two independent experiments; MMVPB: Medicines for Malaria Venture Pathogen Box;
†The chemical classes of the compounds were obtained from PubChem NIH database (https://pubchem.ncbi.nlm.nih.gov).

#### Anti-trypanosomal concentration–response of selected MMVPB

The 70 preselected compounds were tested at serially diluted concentrations to determine their median inhibitory concentrations (IC_50_). Overall, the obtained results showed that compounds displayed moderate to highly potent activities with IC_50_ values varying from 9.78 µM (MMV024311) to 0.0023 µM (MMV688180). The activity data for the 7 reference drugs are shown in Table 1, where the anti-trypanosomatid IC_50_ values ranged from 0.002 µM for MMV000062 (pentamidine-trypanosomiasis) to 1.5 µM for MMV000063 (sitamaquine-leishmaniasis) and the antimalarial IC_50 ranged_ from 1.6 µM for MMV000016 (mefloquine) to 8.4 µM for MMV000023 (primaquine). Promising compounds that were identified in this study and that have previously reported activity against trypanosomatid (25) and against other disease targets (38) are listed in Tables 2 and 3, respectively. Their chemical structures and classes are provided in supplementary information (S1 Table). These inhibitors displayed acceptable cytotoxicity profiles against the African green monkey kidney Vero cell line with selectivity indexes greater than 10 for many.

**Table 3:** *In vitro* antitrypanosomal and cytotoxic activities of the 38 MMVPB compounds with known potency against other diseases.

| Compounds<br>No | MMVPB ID | *IC<sub>50</sub> *T.b. brucei*<br>(μM) | **CC<sub>50</sub> Vero | SI | †Chemical Class | #Reported activity- IC<sub>50</sub>/MIC<br>(μM) | #Target diseases |
| --- | --- | --- | --- | --- | --- | --- | --- |
| 1 | MMV687807 | 0.50±0.04 | 11.±1.30 | 23 | Benzamide | MIC= 2.3 | Tuberculosis |
| 2 | MMV687248 | 0.50±0.07 | >100 | >199 | 1H-Benzimidazol-2-amine | MIC= 9.9 | Tuberculosis |
| 3 | MMV687138 | 0.50±0.01 | >100 | >199 | Benzamide | MIC> 100 | Tuberculosis |
| 4 | MMV495543 | 0.70±0.25 | NT | NT | Benzamide | MIC= 81.9 | Tuberculosis |
| 5 | MMV675996 | 0.80±0.02 | NT | NT | cyclohexanecarboxamide | IC<sub>50</sub>= 43 | Filariasis |
| 6 | MMV688763 | 0.80±0.10 | 8.14±0.01 | 10 | pyridazinone | IC<sub>50</sub>= 3.7 | Schistosomiasis |
| 7 | MMV085210 | 0.90±0.10 | >100 | >107 | Benzenesulfonamide | >2 | Malaria |
| 8 | MMV054312 | 1.29±0.42 | >100 | >77 | pyrroloquinoline | MIC= 0.2 | Tuberculosis |
| 9 | MMV667494 | 1.30±0.50 | >100 | >75 | Quinolone 4-carboxamide | IC<sub>50</sub>= 0.007 | Malaria |
| 10 | MMV024937 | 1.40±0.50 | 14.63±0.60 | 11 | oxazolecarboxamide | IC<sub>50</sub>= 1.2 | Malaria |
| 11 | MMV010576 | 1.50±0.30 | >100 | >66 | 2-amino Pyridines | IC<sub>50</sub>= 0.1 | Malaria |
| 12 | MMV687812 | 1.70±0.10 | 2.66±0.30 | 1.5 | 2-Pyrazinecarboxamide | MIC= 3.5 | Tuberculosis |
| 13 | MMV022029 | 1.70±0.70 | 13.10±0.20 | 8 | biaryl sulfonamide | IC<sub>50</sub>= 0.8 | Malaria |
| 14 | MMV153413 | 1.70±0.10 | >100 | >57 | tetrasubstituted thiophene | MIC< 0.25 | Tuberculosis |
| 15 | MMV687703 | 1.70±0.06 | >100 | >56 | Benzimidazole | MIC= 38.5 | Tuberculosis |
| 16 | MMV062221 | 1.70±0.60 | NT | NT | phenylpyrazolamine | IC<sub>50</sub>>2 | Malaria |
| 17 | MMV022478 | 1.70±0.01 | 11.42±0.07 | 6 | pyrazolo (1.5-a)pyrimidine | IC<sub>50</sub>= 0.8 | Malaria |
| 18 | MMV028694 | 1.80±0.10 | 10.36±0.10 | 6 | 2.4 disubstituted pyrimidine | IC<sub>50</sub>= 1.1 | Malaria |
| 19 | MMV024035 | 1.80±0.10 | 35.75±0.08 | 19 | Thiophene carboxamide | IC<sub>50</sub>= 0.6 | Malaria |
| 20 | MMV676512 | 1.90±0.07 | NT | NT | 1H-Imidazole-5-carboxamide | MIC= 82.7 | Tuberculosis |
| 21 | MMV661713 | 1.90±0.02 | >100 | >51 | 4-pyridyl-2-aryl pyrimidine | MIC= 12.5 | Tuberculosis |
| 22 | MMV687251 | 1.90±0.01 | 6.4±2.0 | 3 | pyrimidine | MIC= 15.4 | Tuberculosis |
| 23 | MMV020670 | 1.90±0.02 | NT | NT | 6-naphthyridine-2-carboxamide | IC<sub>50</sub>= 1.1 | Malaria |
| 24 | MMV687765 | 2.30±0.30 | >100 | >44 | Pyrimidine | MIC= 8.9 | Tuberculosis |
| **25** | **\*MMV675968** | **2.80±0.07** | **>100** | **>35.6** | **aminoquinazoline** | **IC<sub>50</sub>= 0.1** | **Cryptosporidiosis** |
| 26 | MMV688124 | 2.90±0.08 | >100 | >34 | Benzenesulfonamide | MIC= 45.6 | Tuberculosis |
| 27 | MMV688703 | 3.18±1.30 | NT | NT | Pyridines | IC<sub>50</sub>= 99 | Toxoplasmosis |
| 28 | MMV688417 | 4.38±0.90 | 13.60±2.90 | 3 | Pyrazolo [3,4-d]pyrimidinamine | IC<sub>50</sub>= 100 | Toxoplasmosis |
| 29 | MMV023969 | 8.09±0.30 | 26.26±0.40 | 3 | Isoquinoline | MIC= 11.8 | Tuberculosis |
| 30 | MMV688761 | 8.33±0.09 | >100 |  | Benzamide | IC<sub>50</sub>= 0.3 | Schistosomiasis |
| 31 | MMV688768 | 8.36±0.08 | >100 |  | 2,3 disubstituted indole | IC<sub>50</sub>= 3.8 | schistosomiasis |
| 32 | MMV023233 | 8.43±0.06 | 29.75±0.70 | 3 | quinolineamine | IC<sub>50</sub>= 0.2 | Malaria |
| 33 | MMV006901 | 8.45±0.05 | 14.60±0.02 | 1,73 | 2,4-aminoquinoline | IC<sub>50</sub>= 0.6 | Malaria |
| 34 | MMV688854 | 8.50±0.06 | 13.41±1.50 | 1,57 | pyrazolo pyrimidineamine | IC<sub>50</sub>= 0.2 | Cryptosporidiosis |
| 35 | MMV016136 | 8.61±0.30 | NT | NT | pyrazolo pyridineamine | IC<sub>50</sub>= 0.6 | Malaria |
| 36 | MMV011511 | 8.94±0.70 | >100 | >11 | piperidineamine | IC<sub>50</sub>= 1.2 | Malaria |
| 37 | MMV676411 | 9.44±0.09 | NT | NT | Propanamide | MIC= 91.3 | Tuberculosis |
| 38 | MMV024311 | 9.78±0.20 | NT | NT | 1H-Indole | MIC= 19.5 | Tuberculosis |

\* MMVPB compounds were tested in culture against *T. b. brucei* at serially diluted concentrations. Median inhibitory concentrations (IC<sub>50</sub>) were generated from concentration–response curves using GraphPad Prism 8.0 software.
\*\*Cytotoxicity of inhibitors toward Vero cells was assessed using the resazurin viability assay. Selectivity indexes (SI) were calculated as CC<sub>50</sub> (Vero cells)/IC<sub>50</sub> (*T.b.brucei*).
The results are means±standard deviation (SD) from duplicate values obtained from two independent experiments; MMVPB: Medicines for Malaria Venture Pathogen Box;
†The chemical classes of the compounds were obtained from PubChem NIH database (https://pubchem.ncbi.nlm.nih.gov).

Examination of the 38 antitrypanosomal hits listed in Table 3 indicates that they belong to 18 different chemical classes, *viz*. Benzimidazole, Benzamide, Pyrimidine, Pyrazolopyrimidine, Thiophene, Carboxamide, Imidazole, Quinolone, Oxazole, Naphthyridine, Pyrazine, Benzene, Biaryl, Pyridines, Thiophene, Amino Quinazoline, Phenyl Pyrazolamine and Pyridazinone. In the framework of our quest for trypanosomal DHFR inhibitors, we have opted to interrogate the diaminoquinazoline derivative, MMV675968 (2,4 diaminoquinazoline), that was identified with an antitrypanosomal IC_50_ of 2.8 µM and a selectivity index >35. Of note, the literature indicates that the diaminoquinazoline core is a suitable ligand for protein inhibition, including parasite enzymatic targets [22, 23]. More specifically, the quinazoline core was reported as a good motif for the inhibition of trypanothione reductase (TR), a validated therapeutic target for antitrypanosomal drug development [24, 25]. Moreover, the MMV675968 hit compound was previously reported to inhibit the dihydrofolate reductase (DHFR) enzyme both in *Cryptosporidium* and in trypanosomes [26–28]. Other supporting facts for this choice are the diaminoquinazoline analogs trimetrexate and methotrexate, which were reported by Gibson et al. [29] to possess picomolar Ki values against DHFR. Based on this rationale, we hypothesize that this core structure has strong therapeutic potential and could therefore serve as a good starting point for the discovery of novel structural pharmacophores capable of inhibiting vital enzymes (DHFR and TR) of *Trypanosoma brucei brucei* in view of developing new antitrypanosomal drugs. In this line, 23 analogs of MMV675968 readily available at MMV were also tested for antitrypanosomal and cytotoxicity activities.

#### Rudimentary Structure-Activity-Relationship (SAR) study with 23 analogs of 2,4-diaminoquinazoline (MMV675968)

Based on the antitrypanosomal inhibitory potency and selectivity (IC_50_ 2.8 µM; SI >35) of compound 2,4-diaminoquinazoline and its inhibition of enzymatic targets of pharmacological potential, 23 analogs were available and donated by MMV for testing. The results achieved against *T. b. brucei* and Vero cells are summarized in **Table 4**.

**Table 4:**
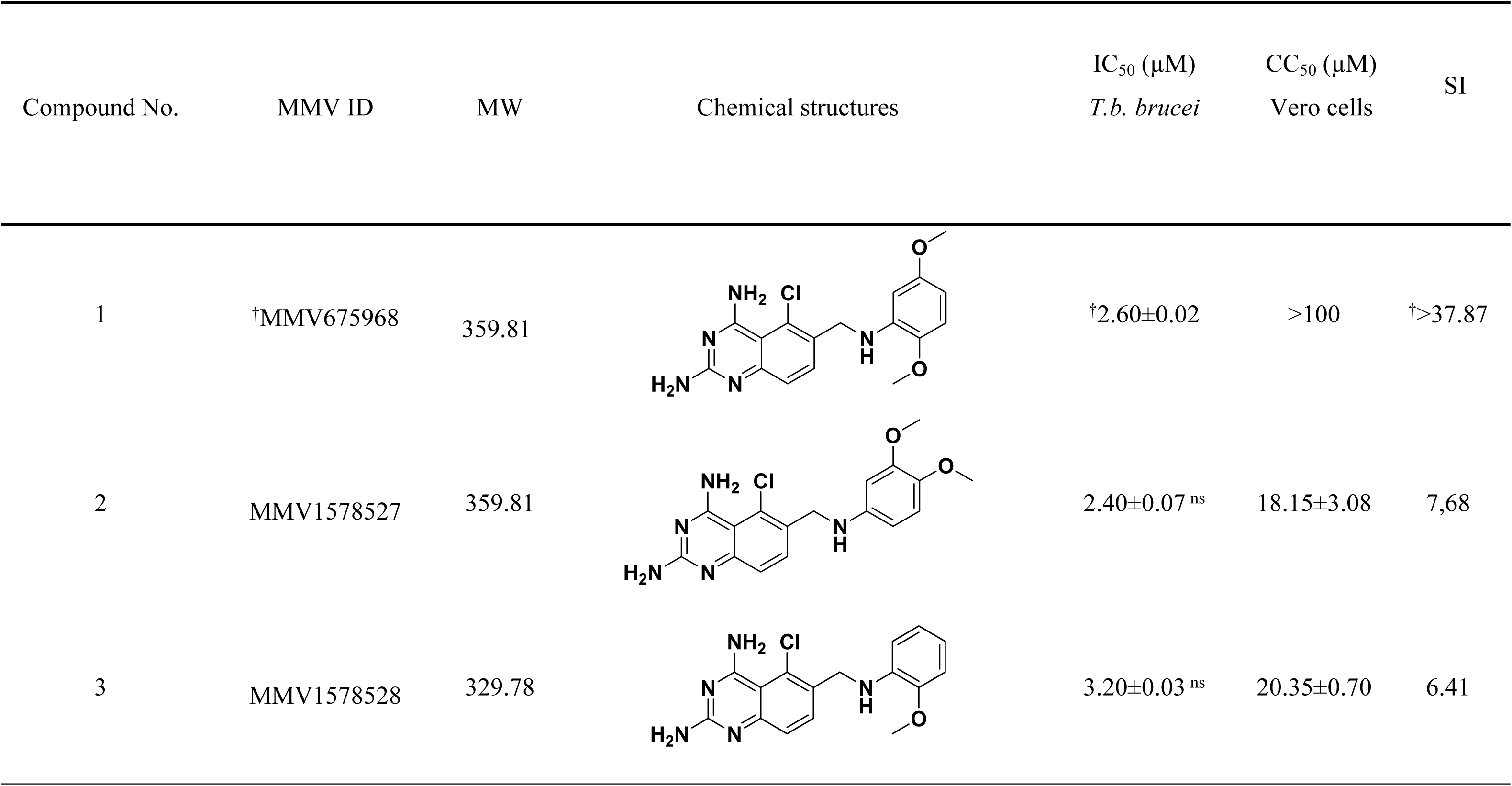
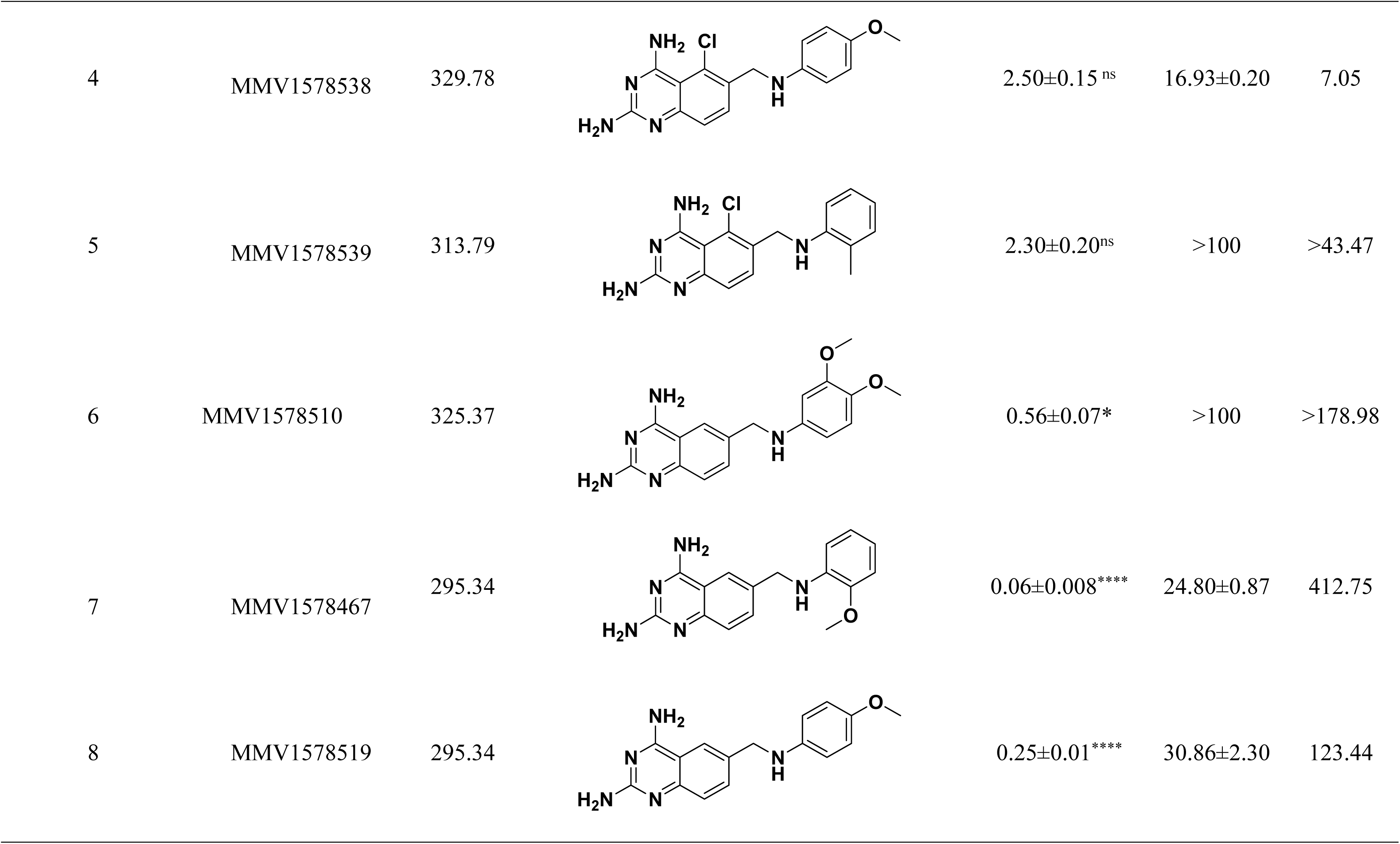
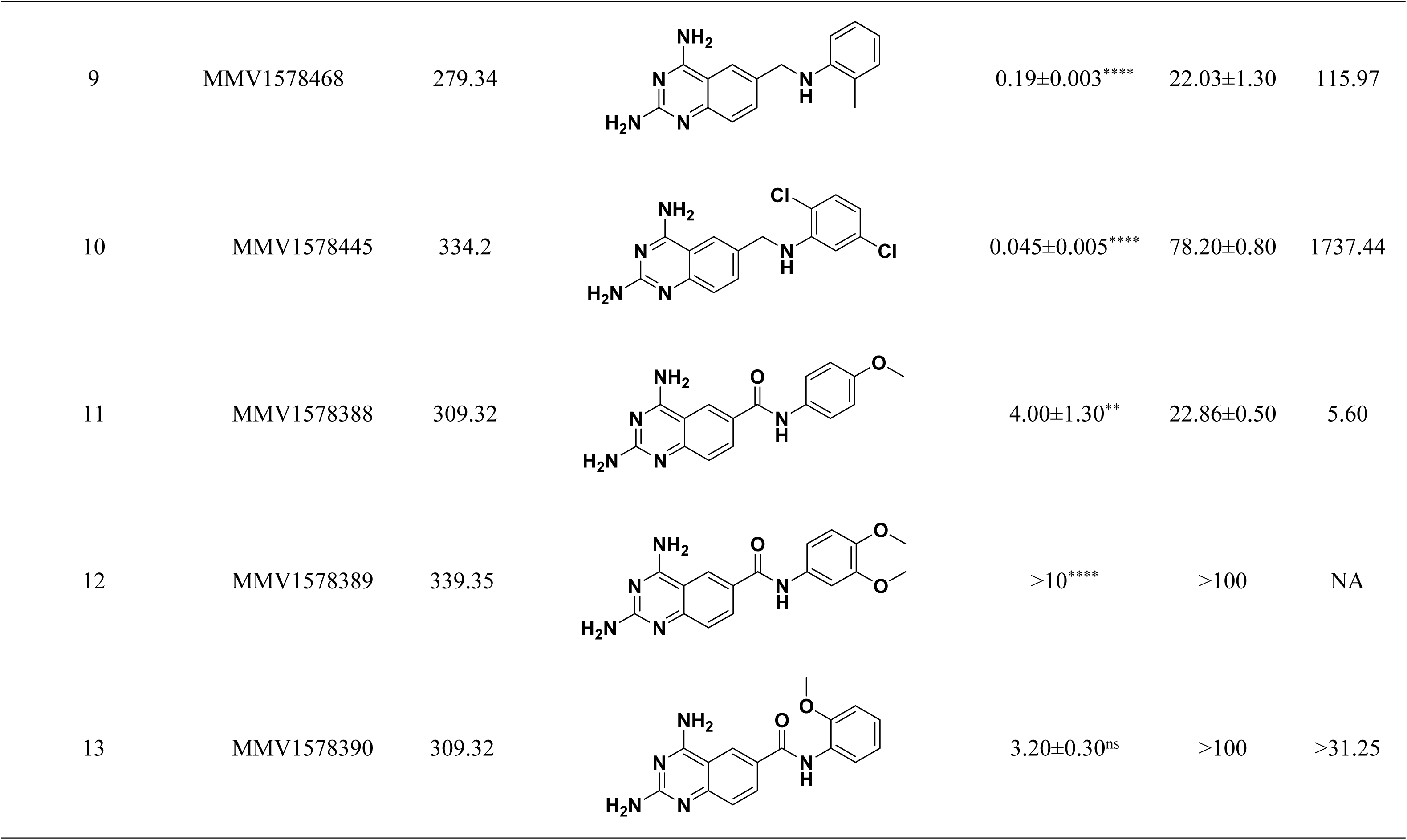
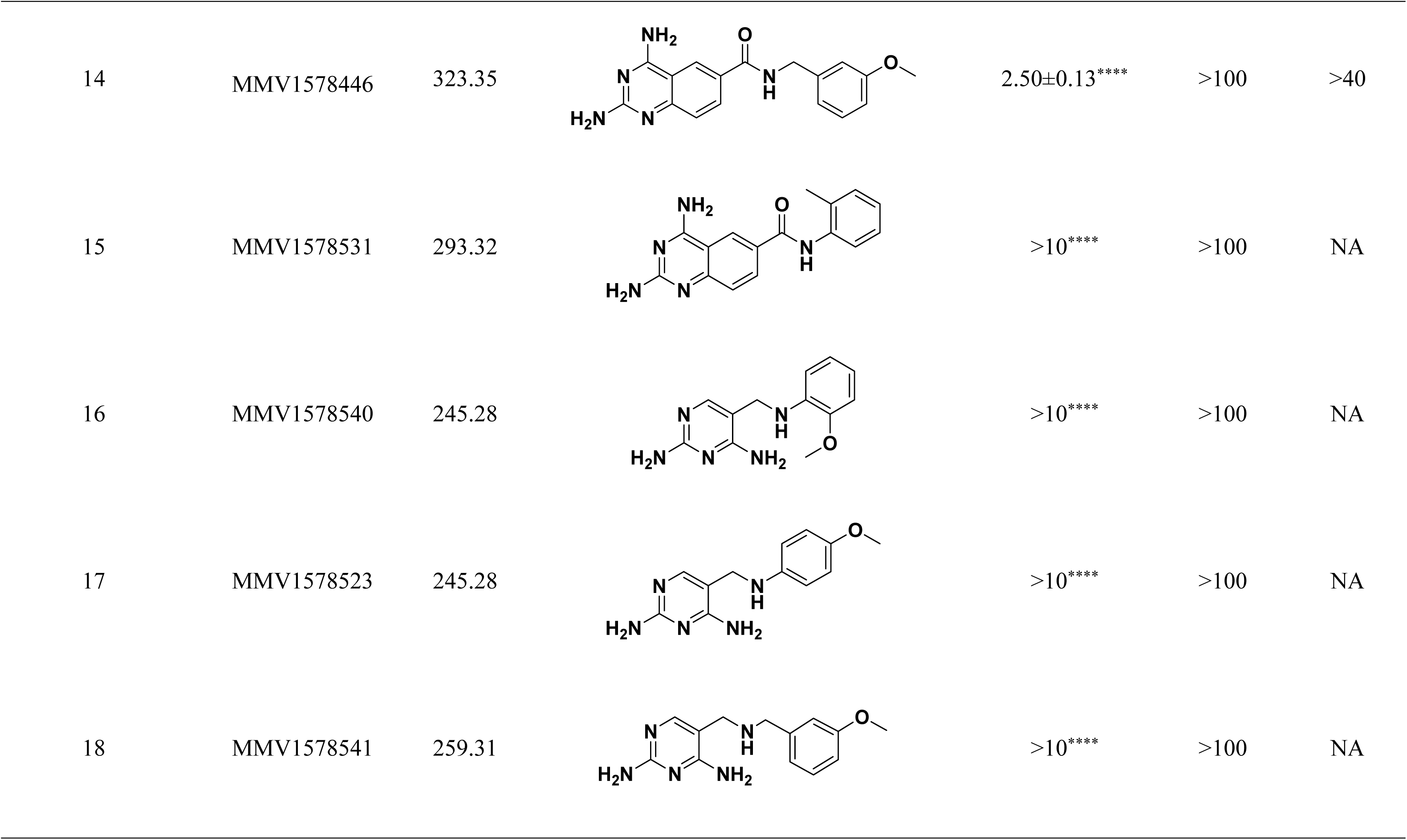
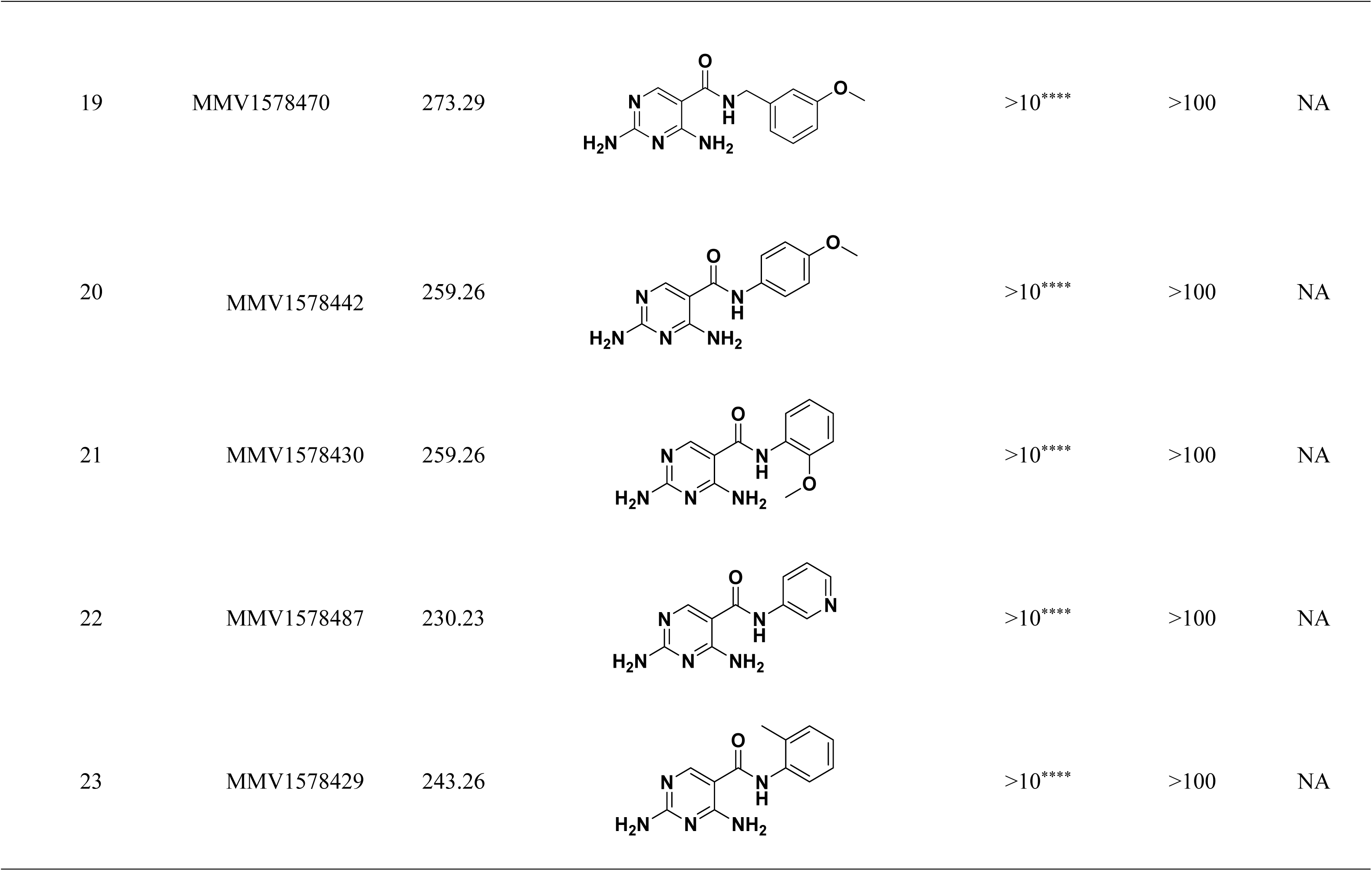
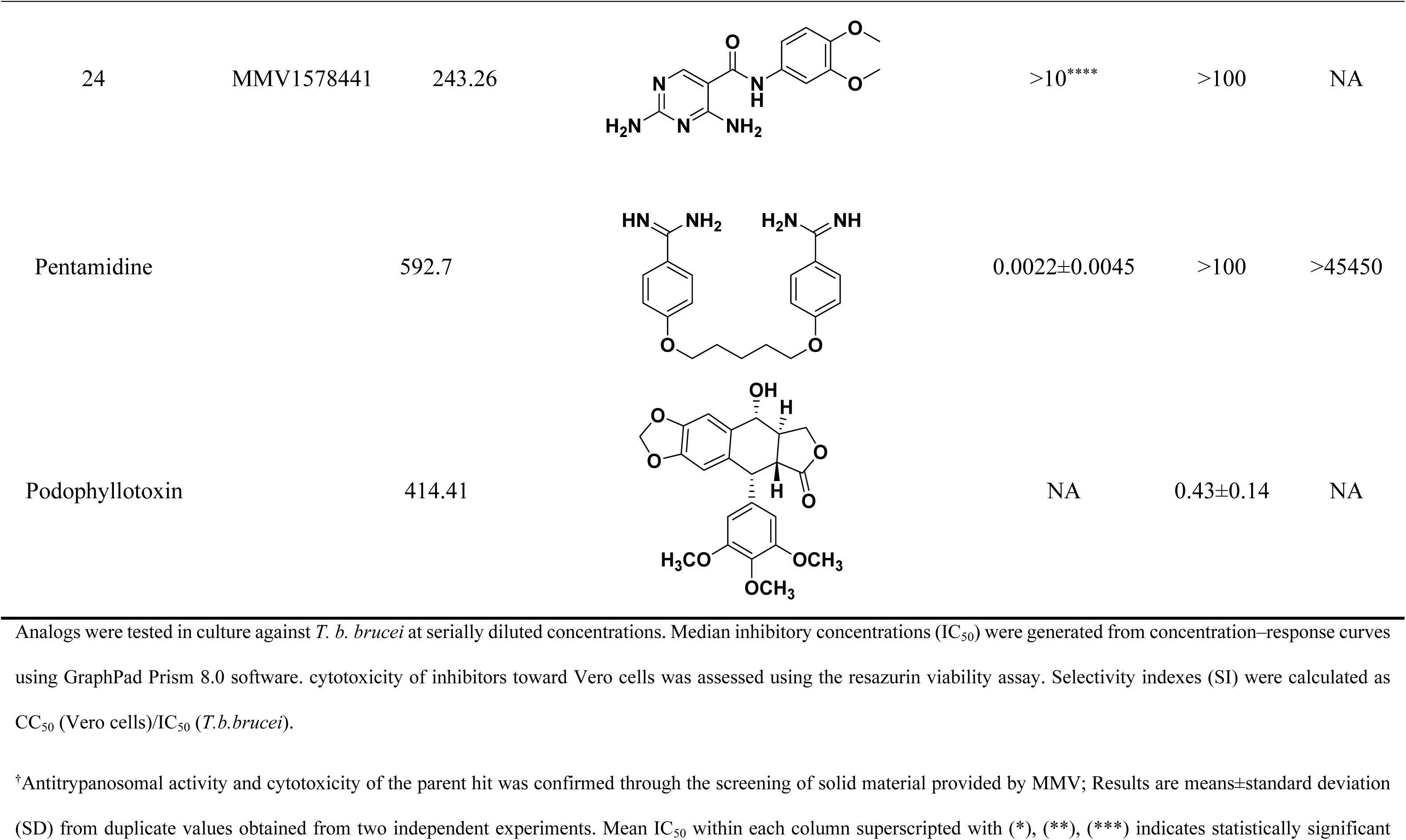
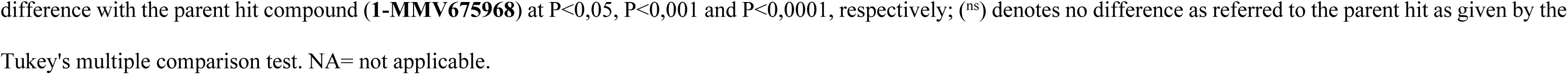
Rudimentary SAR study of 5-chloro-2,4-diaminoquinazoline (MMV675968) analogs for improved antitrypanosomal activity and selectivity.

From the analysis of the results, analogs **2-5** bearing different substitution patterns around the phenyl moiety of hit compound **1** while keeping unchanged the 5-chloro-2,4-diaminoquinazoline core displayed an average IC_50_ value of 2.6 µM, similar to that of parent compound **1** (IC_50_ 2.6-2.8 µM). Similarly, an indifference or even a loss of activity was observed for analogs **11-15** (IC_50_ 2.5 to ˃10 µM), following the incorporation of an amide functionality in analogs **2**, **3**, **4**, **5**, suggesting that neither the substitution pattern of the phenyl nor the amide group are essential for the antitrypanosomal activity. Additionally, removal of the chlorine atom from the core moiety resulted in a 4-(compound **6**) to 53-fold (compound **7**) increase in antitrypanosomal potency (**series 6-9**) with respect to their corresponding congeners **3** and **2 (Series 1-5)**. In contrast, the introduction of chlorine atoms not on the 2,4-diaminoquinazoline core moiety but at positions 2 and 5 of the phenyl group afforded the most active and selective compound identified in this study, analog **10** (IC_50_ 0.045 µM; SI 1737). Unfortunately, the replacement of the 2,4-diaminoquinazoline core by the 2,4-diaminopyrimidine core moiety led to a total loss of activity (series **16-24**, IC_50_˃10 µM).

Overall, SAR studies on the parent hit **1** (MMV675968) (IC_50_ 2.6-2.8 µM; SI ˃37) allowed the improvement of the *in vitro* potency of analogs against bloodstream forms of *Trypanosoma brucei brucei* with up to ∼58-fold increase in activity with analogs **7** and **10** bearing the higher promise (IC_50_ 0.06 and 0.045 µM and SI 412 and 1737, respectively) (**Table 4**). In addition, analogs **7** and **10** duly fitted the lead-like properties, as stipulated in Lipinski’s rule of five (**Fig 2**).

**Fig 2.**
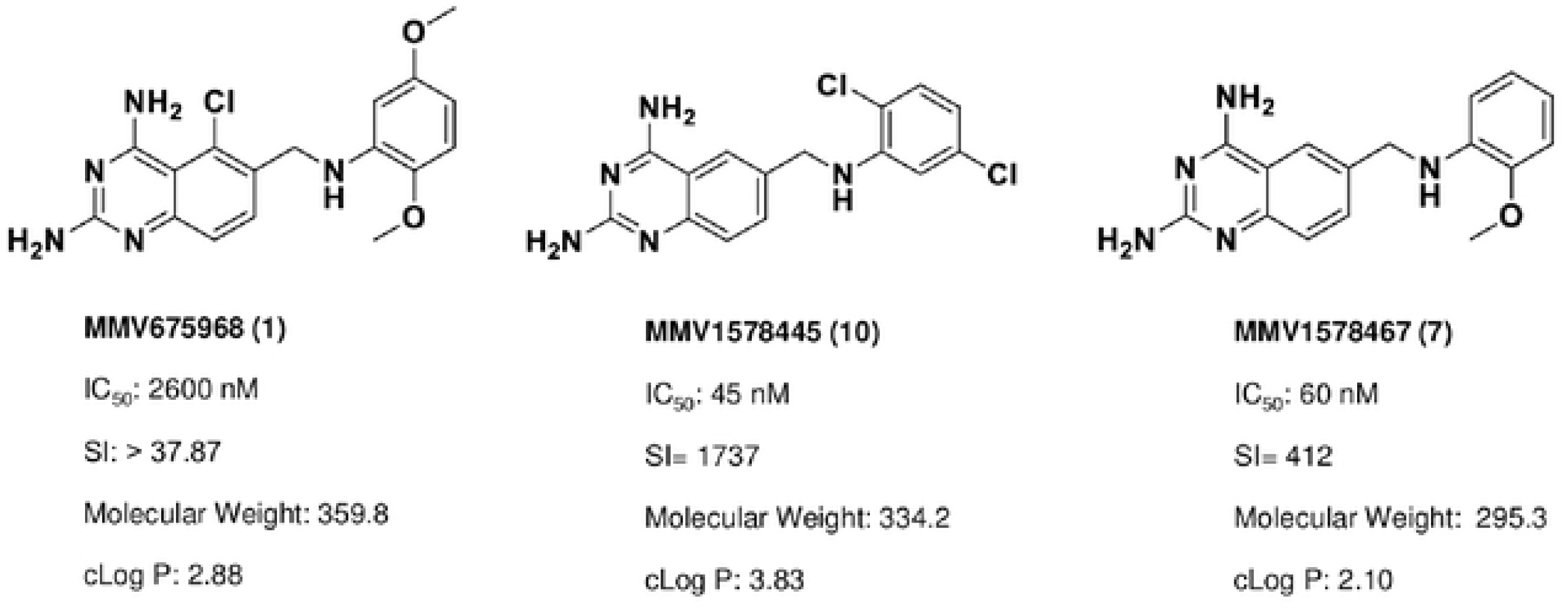
Physicochemical properties of the parent hit (MMV675968) and the prioritized analogs (7, 10). The prioritized analogs adhered to Lipinski’s rule of five (MW ≤ 500 Da, N or O ≤10; NH or OH ≤5, clogP ≤5) [30].

### *In silico* exploration of trypanosomal DHFR and TR inhibition by analogs 7 and 10

#### Accessible protonation states of the compounds and molecular docking

While molecules containing ionizable groups such as amines and carboxylates are stored in databases as neutral entities, they are mostly ionic under physiological conditions. For instance, amines become protonated to the quaternary form while carboxyl and other acidic groups such as phosphates and sulphates or hydroxylamines are deprotonated [31, 32]. This has implications on *in silico* screening experiments as the protonation state tends to influence the binding strength and pose of a ligand within a binding pocket [31, 33]. To account for the protonation states of compounds **7** and **10** at physiological pH, the dimorphite-DL [33] program and MolGpka [34] webserver were utilized as detailed in the methods section. This revealed that positions 2 and 4 of the 2,4 – diaminoquinazoline moiety of both compounds were ionizable at physiological pH [compound **7**: position 2 (pKa = 7.1) position 4 (pKa = 6.5); compound **10**: position 2 (pKa = 7.0), position 4 (pKa = 6.5)]. Hence, a total of four states were considered for each of the compounds – unprotonated (cmpd_7_unprot, cmpd_10_unprot), protonated at position 2 (cmpd_7_prot_2, cmpd_10_prot_2), protonated at position 4 (cmpd_7_prot_4, cmpd_10_prot_4) and protonated at both positions – double protonated (cmpd_7_prot_2&4, cmpd_10_prot_2&4).

Post docking analyses revealed that, in DHFR, all the accessible protonation states of both compounds shared similar poses within the same ligand recognition site in the binding pocket. This resulted in hydrogen bond formation between the diaminopyrimidine moiety and key DHFR ligand recognition residues [35–37], including the highly conserved Asp54, and Val32 (**Figs 3A and 3B**). Unlike DHFR, the TR enzyme consists of a wide binding site, where substrates and inhibitors have been shown to adopt different conformations with stacking observed for some inhibitor binding poses [38, 39]. This explains the relatively diverse manner in which the different protonation states of compounds **7** and **10**, docked within the binding pocket (**Figs 3C and 3D**).

**Fig 3.**
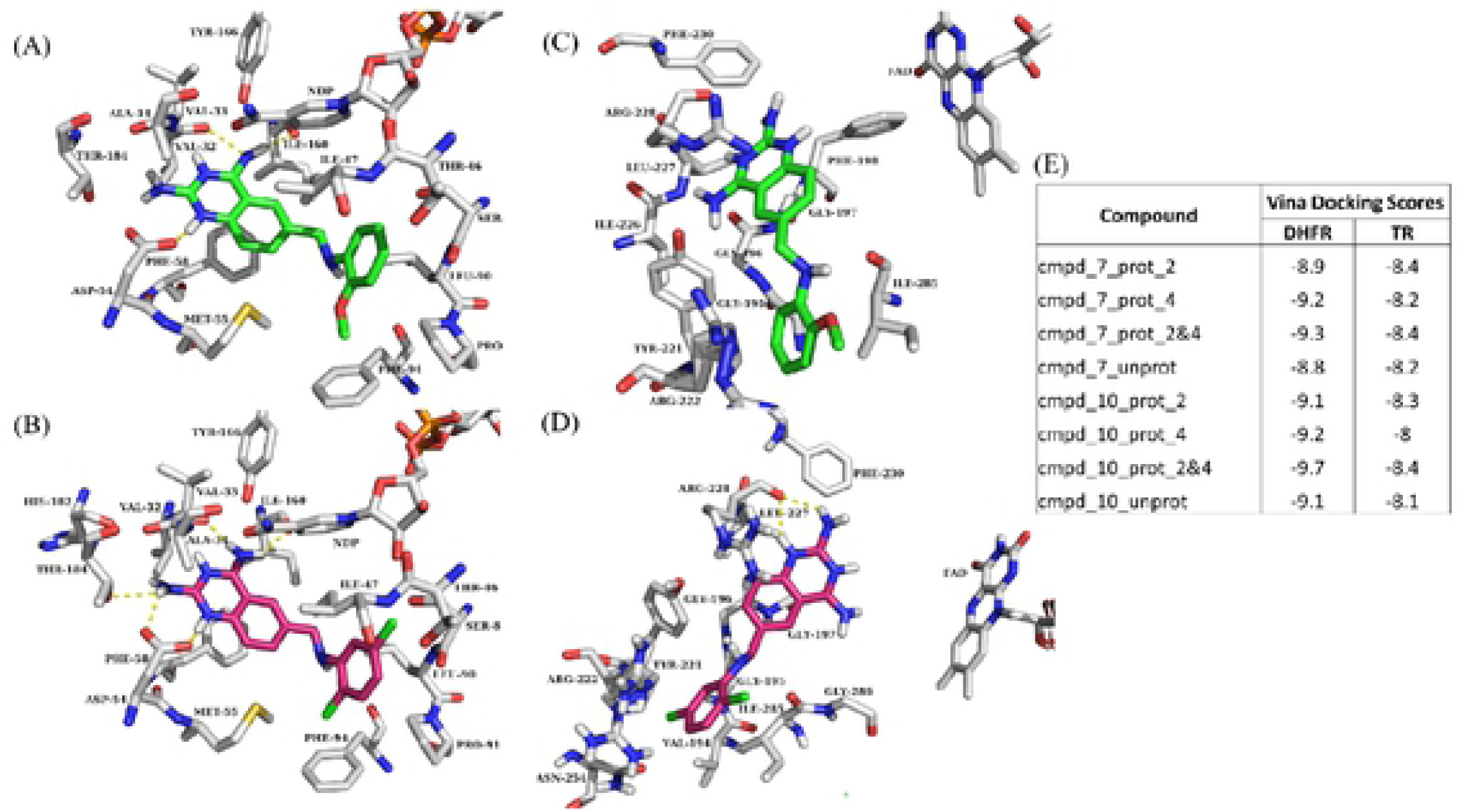
Docking poses and vina docking scores. (A) Docking pose of double protonated compound **7** (cmpd_7_prot_2&4) in DHFR. (B) Docking pose of cmpd_7_prot_2&4 in TR. (C) Docking pose of double protonated compound **10** (cmpd_10_prot_2&4) in DHFR. (D) Docking pose of cmpd_10_prot_2&4 in TR. (E) Table showing vina docking scores for all the protonation states of compounds **7** and **10**. Compound **7** is colored in green while compound **10** is colored in dark pink. Cmpd_7_prot_2 and cmpd_7_prot_4 stand for compound **7** protonated respectively at position 2 and 4 of the 2,4 – diaminoquinazoline ring, cmpd_7_unprot – unprotonated compound **7**, and cmpd_7_prot_2&4 – double protonated compound **7** (protonated at both positions 2 and 4). The same notations apply to compound **10**.

#### Ligand conformational refinement and binding stability monitoring through molecular dynamics simulations

The dynamicity of drug binding and molecular recognition is only partially accounted for - through ligand flexibility - during docking [40] . Thus, MD simulation of the protein-ligand complex is required to further refine the predicted pose from docking and to ascertain the stability of binding. Here 100 ns of all-atom MD simulations were performed on all the eight predicted poses from docking alongside the holoenzymes, making a total of ten systems (i.e., DHFR and TR bound to cmpd_7_prot_2, cmpd_7_prot_4, cmpd_7_prot_2&4, cmpd_7_unprot, cmpd_10_prot_2, cmpd_10_prot_4, cmpd_10_prot_2&4, cmpd_10_unprot, DHFR_holo and TR_holo). Plots of protein RMSD and Rg were used to check for convergence of the different trajectories between the ligand unbound and bound states.

While the number of hydrogen bonds formed by a ligand within the binding site of an enzyme is crucial for its stability, monitoring the ligand RMSD across the simulation reveals the post docking dynamics of the bound ligand, which informs on the stability and strength of binding of the ligand [36] . To monitor the stability of binding of the different compounds, ligand RMSD with respect to protein structure and hydrogen bond numbers were computed across the different simulations. This revealed stable binding for all the DHFR bound compounds, forming between 1 to 7 hydrogen bonds (**Fig 4**). The total number of hydrogen bonds formed tended to increase with additional protonation, such that the unprotonated form had the lowest maximum number of hydrogen bonds, while the double protonated form had the highest (**Fig 4A**). Unlike the DHFR bound compounds, TR bound compounds generally portrayed reduced stabilities. For instance, cmpd_7_prot_4 and cmpd_10_prot_2 lost hydrogen bonds completely during the simulation, with the later exiting from the binding site (**Fig 4B**). On the other hand, although the double protonated form of compound **7** (cmpd_7_prot_2&4) and cmpd_**10**_prot_4 maintained some hydrogen bonding during the simulation, their RMSD values varied considerably, pointing to unstable binding. It has been shown that microenvironmental differences within the binding site of an enzyme can influence its preference for binding to certain protonation states of a compound [41]. Thus, it is possible that while compounds **7** and **10** are good binders of DHFR in all their accessible protonation states, both compounds may have difficulties binding to and staying within the binding site of the TR enzyme, with only their unprotonated forms, as well as the position 2 protonated compound **7**, and double protonated compound **10** having the possibility of binding successfully to the enzyme.

**Fig 4.**
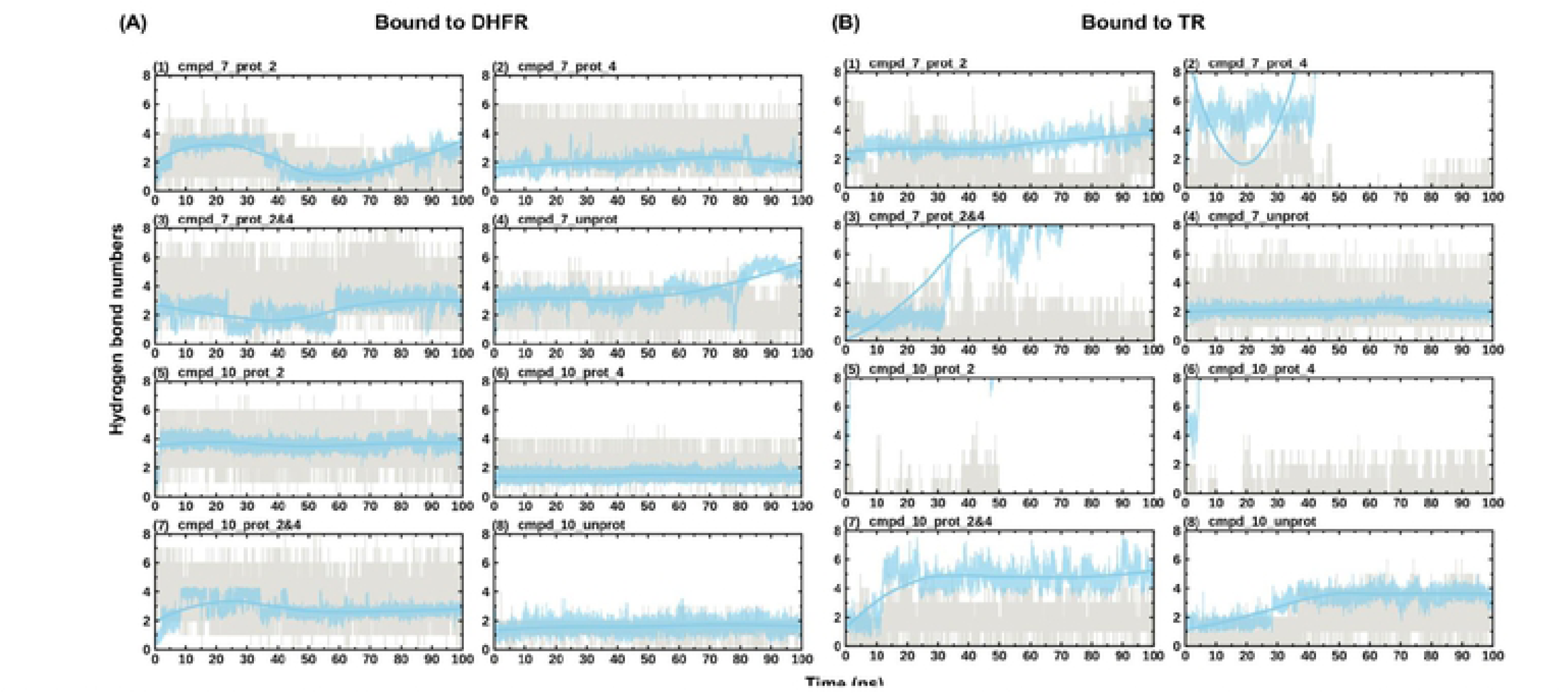
Evolution of ligand RMSD with respect to protein and hydrogen bond numbers during simulation. (A) Shows the evolution of the different protonation states of the compounds in DHFR. (B) Shows the evolution of the different protonation states of the compounds in TR. Hydrogen bond numbers are shown in grey while the evolution of the ligand RMSD is shown in blue. RMSD values have been scaled up by a factor of 10 to fit into the scale of the hydrogen bonding numbers plot.

To further elucidate the key residues responsible for ligand recognition in both enzymes, ligand clustering of the last 10 ns of the MD trajectories was conducted as outlined in the methods section. In DHFR, all the ligands populated a single cluster, further supporting the stability of binding of both compounds to DHFR in all their protonation states. This, however, was not the case with TR, where multiple clusters were populated particularly for the double protonated compound **7**, and the single protonated forms of compound **10**. Examination of the ligand interaction patterns of the representative structures from the populated single clusters revealed the following residues Val32, Asp54, Ile160 as key interacting residues, forming hydrogen bonds with the diaminoquinazoline (DMQ) moiety of both compounds in DHFR (**Table 5**). The rest of the residues mainly formed stacking and other interactions. In TR however, no key residues were seen cutting across all the systems in terms of interaction. This may be expected due to the wide nature of the TR binding pocket as explained above.

**Table 5:** Residues interacting with compounds 7 and 10 within the binding sites of DHFR and TR.

| DHFR | Ring 1 of DMQ | Ring 2 of DMQ | tail ring |
| --- | --- | --- | --- |
| cmpd_7_prot_2 | Val32, Ile160, NDP | Ile47, Phe58 | Pro91, Met55 |
| cmpd_7_prot_4 | Asp54, Ile160 | Phe58 | Pro91, Leu90 |
| cmpd_7_prot_2&4 | Asp54, Val32, NDP | Phe58 |  |
| cmpd_7_unprot | Val32, Asp54, Ile160 | Val33, Ala34, Ile47, NDP | NDP |
| cmpd_10_prot_2 | Val33, Asp54 | Ile160, Phe58, NDP | Leu90, Pro91 |
| cmpd_10_prot_4 | Val32, Asp54, Ile160 | Phe58, NDP | Ile47, Pro91, Phe94 |
| cmpd_10_prot_2&4 | Val32, Asp54, Ile160, | Phe58 | Met55, Leu90, Pro91, Phe94 |
| cmpd_10_unprot | Asp54, Met55, Ile160, NDP | Phe58, NDP | Ile47, Pro90, Phe94 |
| TR |  |  |  |
| cmpd_7_prot_2 | Tyr221, Asn223, Arg228 |  | Arg222, Ile285 |
| cmpd_7_unprot | Arg228 | Gly196 | Arg222, Ile285 |
| cmpd_10_prot_2&4 | Gln165, Arg287, Lys306 |  | Pro167, Tyr221, Arg222, Ile285 |
| cmpd_10_unprot | Ala284, Ala286, Ala287 | Ile199, Phe198, FAD | Leu332, Met333, Leu334 |

DMQ - diaminoquinazoline

### Tentative elucidation of other modes of action of analogs 7 and 10 against *T.b. brucei*

Based on their *in vitro* activity and selectivity profiles and their suitable drug-likeness, compounds **7** and **10** were chosen for further study of their tentative mode of action against *T.b. brucei*.

#### *In vitro* kinetics of *T.b. brucei* killing upon treatment with analogs 7 and 10

Analogs 7 and 10 were assessed at their respective IC_99_, IC_90_, IC_50_ and IC_10_ for their impact on the growth rate of *Trypanosoma brucei brucei* in culture during a complete trypanosome life cycle (72 h) (**Fig 3**). The results indicated a significant concentration-dependent reduction in trypanosome growth (Fig 5A and 5B) compared to untreated cells when exposed to compounds 7 and 10 at their IC_99_, IC_90_, IC_50_ and IC_10_. A similar trend was observed for the positive control (pentamidine-Fig 5C). A rather mild effect of inhibitors on parasite growth was observed at their IC_50_ and IC_10_. Of particular interest, compound 10, similar to the control (pentamidine), completely suppressed the growth of trypanosomes throughout the 72h life cycle, whereas the effect of compound 7 tended to decrease after 60 h of treatment (Fig 5B).

**Fig 5.**
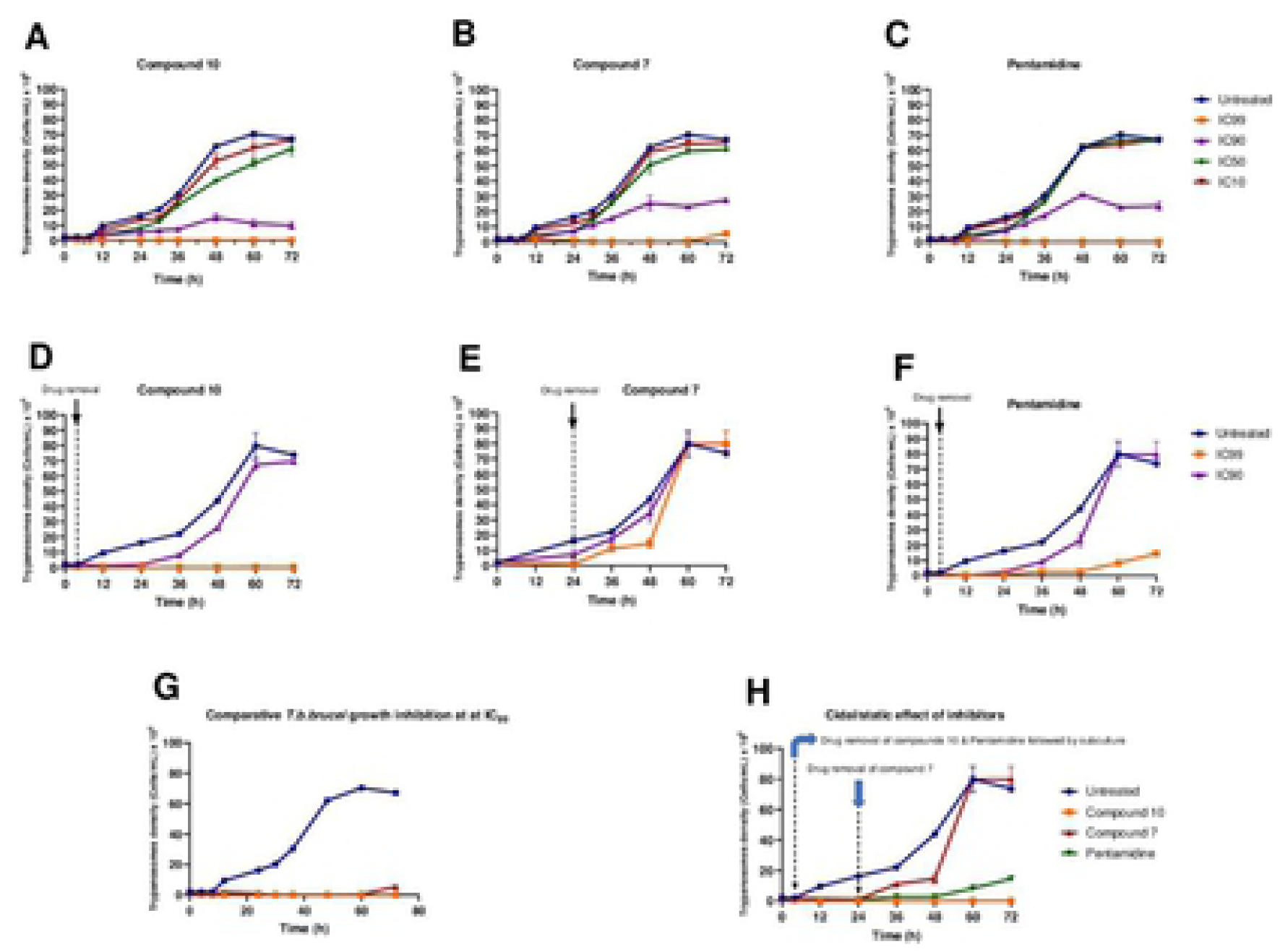
Growth curves of *T.b. brucei* in the presence or absence of various concentrations of compounds 7 and 10 and pentamidine. Parasites were seeded at 2 x 10^5^ cells/mL and incubated for 72 h at 37°C and 5% CO_2_. (Fig 5A-5C) At the indicated time points, trypanosomes were counted microscopically (for IC_10_, IC_50_, IC_90_ and IC_99_ wells), and the obtained cell counts were averaged and then plotted versus time using GraphPad Prism 8.0. (Figs 5D-5F) After treatment for 4 h (analog 10) and 24 h (analog 7), drugs were washed out from the cultures, and the parasites were subcultured in drug-free complete medium and for an additional 68 h and 48 h for analogs 10 and 7, respectively. Cells were then microscopically counted, and the counts were used to plot the growth curves over time. The activity profiles of compounds 7 and 10 and pentamidine are compared in Figs 5G and 5H. The results are expressed as the mean±SD from duplicate experiments.

Inhibitor removal at 4 h for analog **10** and 24 for analog **7** when tested at their respective IC_90_ and IC_99_ followed by subculture in inhibitor-free complete medium indicated a cidal effect for compound 10 throughout the 72h cycle, as all parasites were seemingly dead (Fig 5D). Conversely, at the time of compound **7** removal, the parasites started growing consistently between 24 and 48 h, followed by exponential growth between 48 and 60 h, denoting a rather static effect (Fig 5E). Of note, pentamidine (positive control) portrayed a static profile because its effect started diminishing increasingly from 48 h to 72 h (Fig 5F). Overall, from the comparison of the individual effects of the 3 inhibitors depicted in Fig 5G and 5H, analog 10 appeared to exhibit a more potent (cidal) effect on trypanosomes than analog 7 and pentamidine.

#### Effect of compounds 7 and 10 on plasma membrane integrity

To evaluate their possible effect on plasma membrane integrity, various concentrations (IC_99_; IC_90_, IC_50_, and IC_10_) of compounds **7** and **10** were incubated with parasites for up to 120 minutes. Plasma membrane disruption was examined using the fluorescent probe SYBR Green, which binds to DNA and follows parasite injury permeability caused by direct or indirect action of the compounds. The results indicated that the parasite membrane conserved its integrity even after 120 min of contact with inhibitors. This unchanged status of the plasma membrane was materialized by a nonsignificant difference in the fluorescence profile of drug-treated versus untreated parasites. Conversely, there was a highly significant difference between the effects of compounds 7 and 10 on the one hand and the positive control (saponin) on the other hand, at p<0.05 (**Fig. 6**), denoting no impairment or a rather mild effect of the test compounds on the plasma membrane permeability of the parasite.

**Fig 6.**
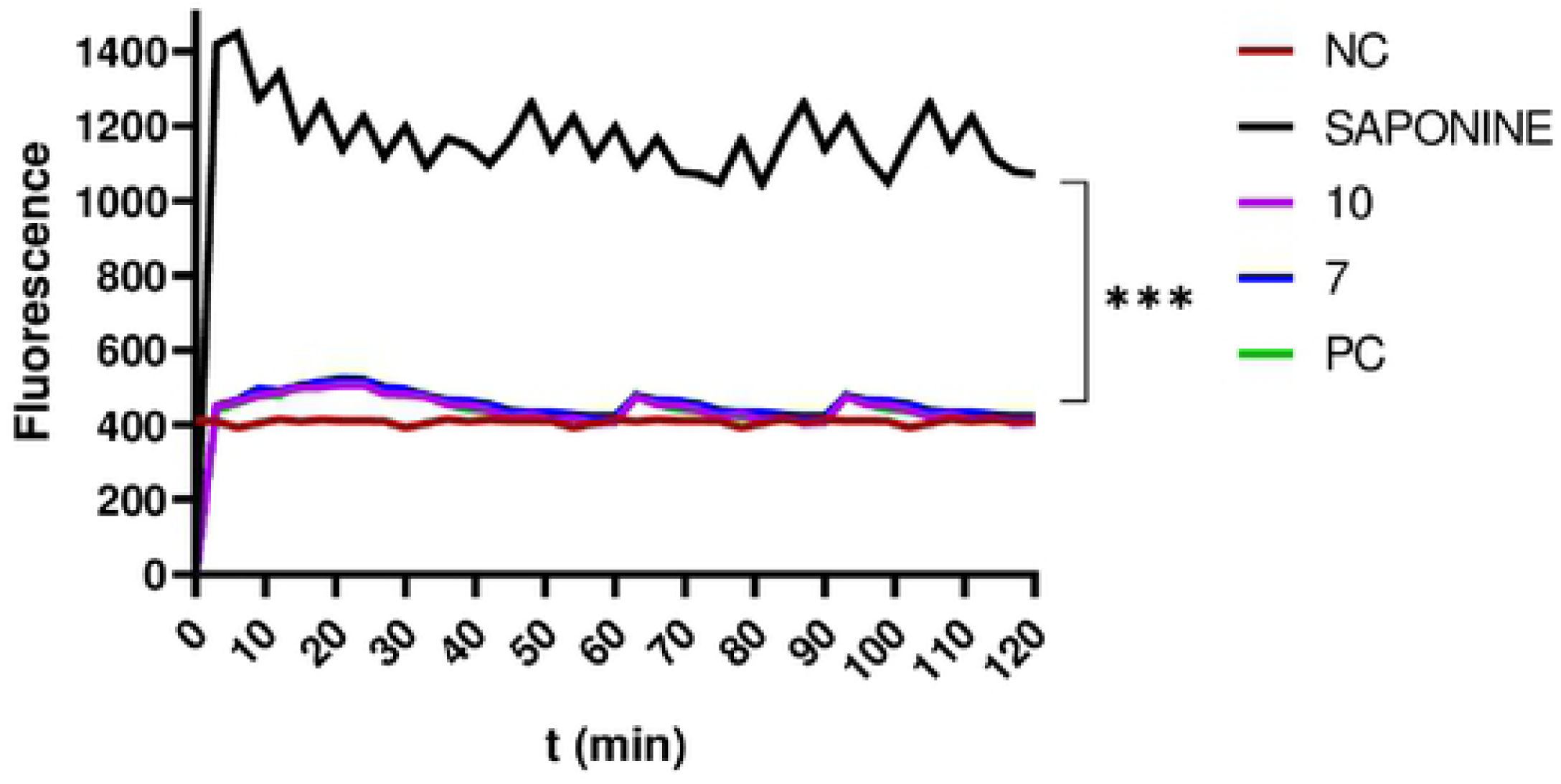
Depiction of the effect of inhibitors on membrane integrity. The fluorescence intensity of trypanosome DNA-SYBR Green (SG) complexion was measured over 120 min. Parasites at 2 x 10^6^ cells/mL were preincubated with SG followed by treatment with compounds **7** and **10** at their IC_99_, IC_90_, IC_50_, and IC_10_ for 120 min. SG uptake was then fluorometrically monitored, and data were obtained to plot corresponding curves versus time. The results were compared to the negative control and the positive control (***Significant difference at P ˂ 0.05). Negative control (NC): trypanosomes untreated with inhibitors; PC: saponin (100% permeability).

#### Induction of oxidative stress in trypanosomes by compounds 7 and 10

We assessed the influence of the test compounds on reactive oxygen species (ROS) production by trypanosomes as an indication of oxidative stress induction. Trypanosomes were incubated with compounds **7** and **10** for 120 min, and the intracellular level of ROS was determined using the fluorescent probe H2DCF-DA. The results obtained indicated no significant change (P˂0.05) in ROS production in treated parasites when compared to the untreated parasites (negative control), contrary to those treated with the positive control (H_2_O_2_) at 0.1% (v/v) (**Fig 7**).

**Fig 7.**
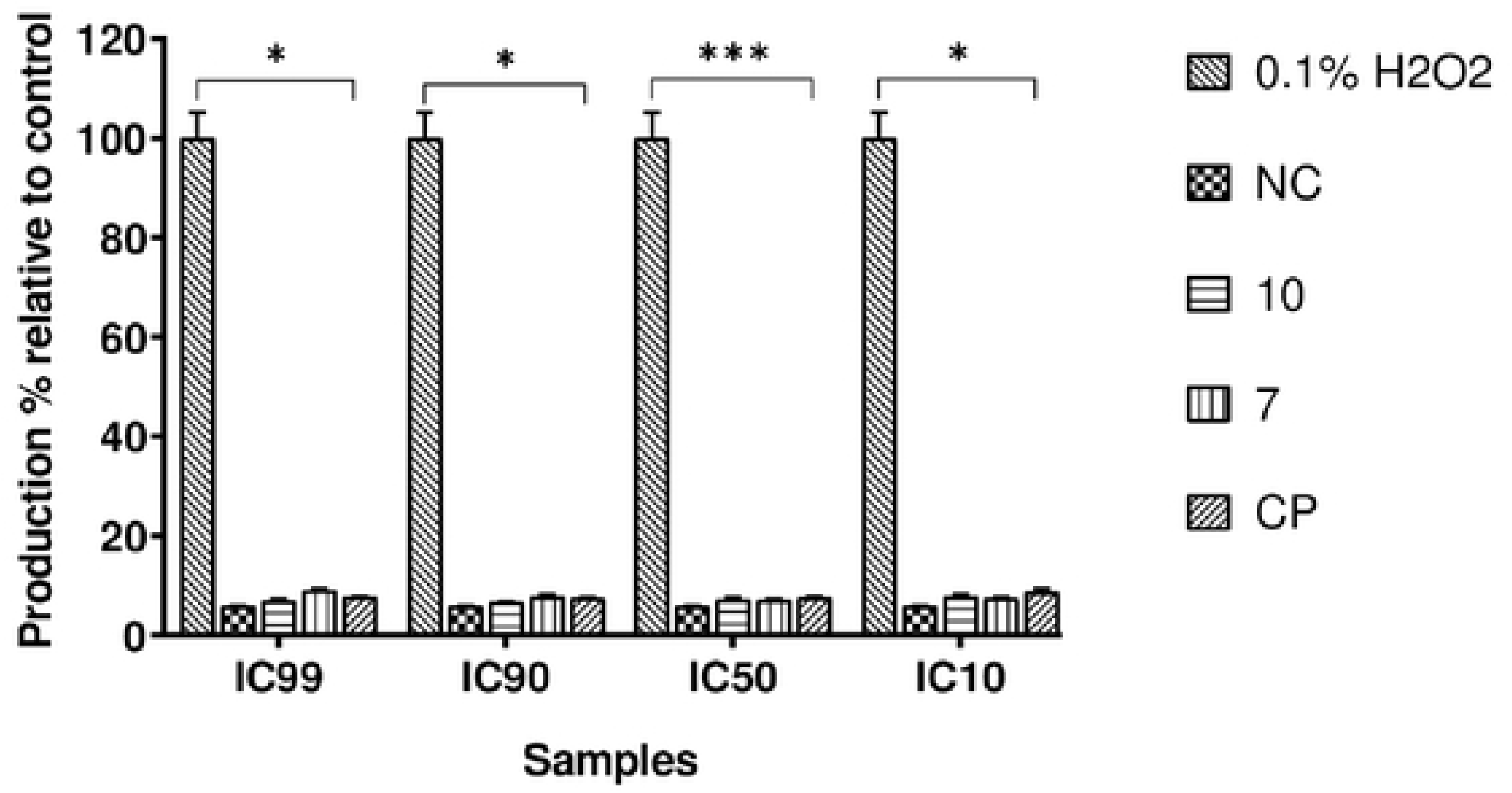
Production of intracellular reactive oxygen species (ROS). Parasites were treated with or without compounds at their respective IC_99_, IC_90_, IC_50_ and IC_10_ for 120 min, and ROS production was measured using DCF-DA reagent. H2O2 (0.1%_)_ was included as a positive control and further used to calculate the ROS production percentages. The obtained results are expressed as the mean±standard deviation of two independent experiments performed in duplicate. * Significant difference at p<0.05.

#### Compound 10 induces DNA fragmentation in trypanosomes as a sign of late apoptosis

To shed light on the mode of parasite killing induced by the inhibitors, we further determined the type of cell death elicited by compounds **7** and **10** using a DNA fragmentation kit. The technique used consisted of immunological detection of BrdU (5’-bromo-2’-deoxyuridine)-labeled DNA fragments into the parasitic cytoplasm for apoptosis and into the culture supernatant for cell-mediated cytotoxicity. The results achieved are summarized in Figure 8 below.

**Fig 8.**
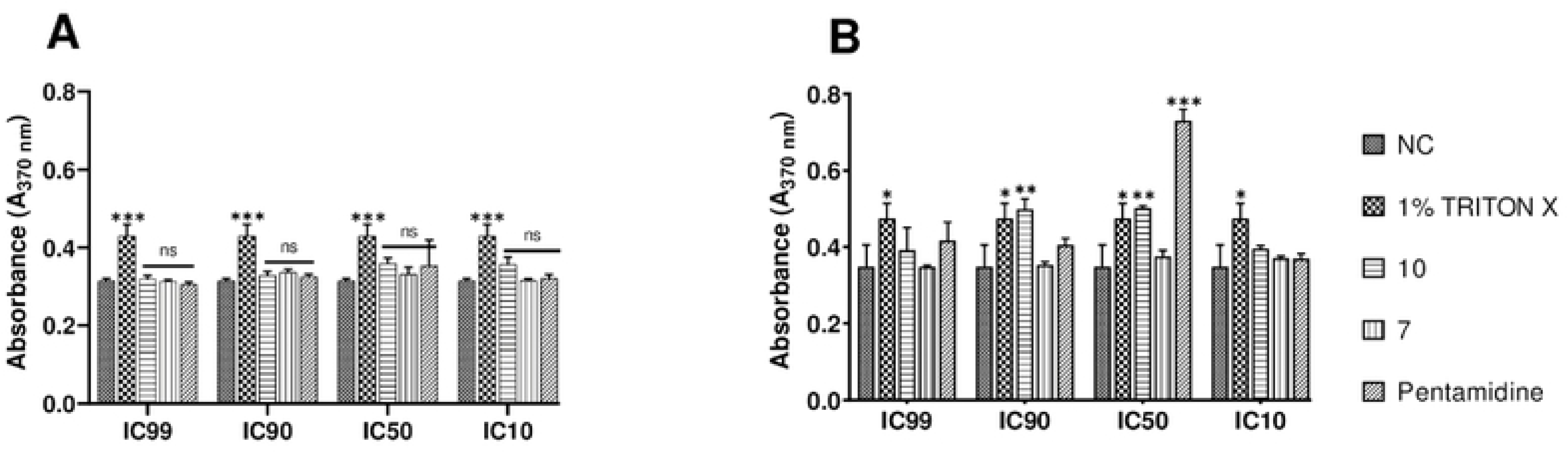
Mediated apoptosis and cytolysis induced by compounds 7 and 10 in bloodstream forms of *Trypanosoma brucei brucei*. BrdU-labeled parasites at 4 x 10^5^ cells/mL were incubated in the presence or absence of compounds **7** and **10** at their IC_99_, IC_90_, IC_50_ and IC_10_ for 4 hours in growth conditions, and the culture supernatant (Fig 8A, cytolysis) and lysate (Fig 8B) were further processed using the Cellular DNA Fragmentation Kit (apoptosis). ***Significant difference at p< 0.001; * Significant difference at p< 0.05; ns: Not significant difference; NC: Negative control (untreated parasites); 1% Triton-X was used as a positive control.

Mediated cell death was depicted through measurement of stronger absorbance emitted by DNA-labeled fragments with a characteristic green color. From the examination of Fig 8A, there was no apparent induction of cell lysis by any of the test compounds, similar to pentamidine relative to Triton-X, which is considered a potent cell disruptor. On the other hand, compound 10 exhibited a significant apoptosis-inducing effect. However, this effect was to a great extent lower than that exhibited by pentamidine, the reference trypanosomiasis drug (**Fig 8B**), at their respective IC_50_s. Of note, mediated apoptosis was not seen for all inhibitors at their IC_99_, IC_90_ and IC_10_, probably because the parasite death rate at IC_99_ and IC_90_ was significantly high to enable the development of the measurable green anti-BrdU-DNA complex, or the parasite killing rate was not significant at IC_10_ (due to subinhibitory concentration) to enable measurement of the drug effect. It should also be pointed out that compound 7 did not exert any effect through the two cell-mediated death modes investigated.

#### Ferric Ion Reducing Antioxidant Power (FRAP) of Inhibitors

To assess the potential of inhibitors as reducing agents, we determined their ferric iron reducing capacity. The results showed that compound **7** exhibited weak Fe^3+^-reducing activity with a median reducing concentration (RC_50_) of 96.37 µM. In addition, compound **10** (RC_50_ ˃ 400 µM) was over ∼ 4-fold less active than compound **7**. The latter showed more potent Fe^3+^-reducing activity than ascorbic acid (165 µM), which was used as a positive control. This finding suggests that compound **7** might exert its antitrypanosomal activity through deprivation of ferric iron bioavailability to trypanosomes.

## Discussion

Sleeping sickness remains a dreadful public health emergency, particularly in endemic regions where it causes significant damage to both cattle and humans [2]. Of sure, control measures have been established for decades and include chemotherapy, which consists of the use of approved drugs. However, these drugs have several drawbacks, such as cumbersome lengthy treatment, adverse and toxic effects, and the development of resistant trypanosome mutants [42, 43]. In addition, following a slackening in control measures, resurgence of the disease was observed in some foci [4]. This situation emphasizes the need for more attention to this neglected tropical disease (NTD) in view of definitely eliminating it. Consequently, new therapeutic options are urgently needed to supply the pipeline of antitrypanosomal drug discovery and development, which might be suitable for optimal management of trypanosomiasis. To achieve this goal, many strategies have been developed to date and have gained successful results. This is the case for drug repositioning, which offers a tangible opportunity to quickly identify new chemical entities with appropriate potency against pharmacologically validated targets in trypanosomes [44].

Within this framework, we screened the Medicines for Malaria Venture’s Open Access Pathogen Box against the bloodstream forms trypomastigotes of *Trypanosoma brucei subsp. brucei*, Lister 427 VSG 221. Out of the 400 compounds tested, 70 were found to inhibit the growth of the parasite by at least 90% (**Fig 1**), and their IC_50_ values ranged from low micromolar (∼9.8 µM) to low nanomolar (IC_50_ ∼ 2 nM). **Veale and Hoppe** [45] obtained approximately similar results from the screening of MMVPB, with approximately 65 hits identified against *T. b. brucei*, although some discrepancies in activity and potency were observed, probably due to differences in the culture and assay conditions used (incubation period, parasite load, etc.). From the 70 compounds that emerged from our preliminary screening, internal controls (reference compounds) were excluded from further studies, including 5 anti-trypanosomatids (MMV637953 (Suramine)-Trypanosomiasis; MMV001499 (Nifurtimox)-Chagas disease; MMV688773 (Benzimidazole)-Chagas disease; MMV000062 (Pentamidine)-Trypanosomiasis; MMV000063 (Sitamaquine)-Leishmaniasis) and 2 antimalarials (MMV000016-Mefloquine and MMV000023-Promaquine). Another set of 25 other compounds previously reported to inhibit trypanosomatid parasites was also identified (Table 2). Selected examples among these included compound **MMV688180,** a benzenesulfonamide (IC_50_ 0.0023 µM) that was the most active against *T. b. brucei* (**Table 2**). This compound was previously reported for its activity toward trypanosomes via the inhibition of N-myristoyl transferase, an enzyme that is essential for the survival and virulence of *T. b. brucei* [46, 47]**(Price et al., 2010; Brand et al., 2012)**. Additionally, **MMV689029** and **MMV689028**, which form part of the benzyl piperazine class of compounds, and the 2,4-substituted furan **MMV688796** were previously reported by **Duffy et al.** [20] as promising starting points for drug discovery against both *T. cruzi* and *T. brucei*. Similarly, the kinase inhibitor **MMV676604** (a 2-aminopyramidine), which exerts trypanocidal activity through inhibition of *Tb*ERK8 (extracellular signal-regulated kinase 8), was reported by **Valenciano et al.** [48]**. MMV652003**, which is a member of the benzamide class of molecules, has also been reported to act on leucyl-tRNA synthetase, a pharmacologically significant target of which is the trypanosome [49, 50]. The butyl sulfanilamide **MMV688467** was proven to inhibit microtubule formation in trypanosomes [51]. The antiplasmodial guanidine **MMV688271** [52] was previously predicted to bind to the DNA minor groove at AT-rich regions [50, 53]. All these compounds were equally excluded from further investigations.

The other set of 38 antitrypanosomal hits identified in this study were previously reported for activity against other disease targets, including malaria (14), tuberculosis (16), toxoplasmosis (2), schistosomiasis (3), cryptosporidiosis (2) and filariasis (1) (**Table 3**). Among these compounds, we opted to investigate the SAR of MMV675968 (2,4 diaminoquinazoline). This option was supported by the generous donation of a small library of 2,4 diaminoquinazoline analogs available at MMV. Of note, the data generated from our work indicated that compound MMV675968 has a favorable profile for further studies, including an antitrypanosomal IC_50_ of 2.8 µM and a selectivity index >35. Interestingly, this hit compound was previously reported to inhibit the dihydrofolate reductase (DHFR) enzyme both in *Cryptosporidium* and in trypanosomes [26–28], where it has been chemically and genetically validated as an antitrypanosomal drug target [54]. Other supporting facts for this choice are the diaminoquinazoline analogs trimetrexate and methotrexate, which were reported by **Gibson et al.** [55] to possess picomolar Ki values against DHFR. Other valuable evidence includes the potential of diaminoquinazoline derivatives against other vital enzymatic targets in trypanosomes, such as the trypanothione reductase enzyme (TR) [24, 25]. In addition, this class of compounds was identified as showing promise from a high-throughput *in vitro* screening of ∼100 000 compounds against *Trypanosoma cruzi* [56], which was further correlated by screening data against *T. brucei* and TR. In addition to their inhibitory activity on the two enzymes mentioned above (TR and DHFR), quinazoline analogs were also identified as potential inhibitors of *Trypanosoma brucei* alternative oxidase (AOT), an important metabolic enzyme. Finally, owing to the broad-spectrum pharmacological properties of quinazoline derivatives (antiparasitic, antiviral, antidiabetic, antibacterial and antioxidant) [57], this scaffold qualifies as a structure of great interest for the identification and development of new drug candidates. More recently, compound MMV675968 emerged as the most active pathogen box compound from a screening against *Toxoplasma gondii* (IC_50_ 0.02 µM; SI 275) [19]. Additionally, it has shown potency against other pathogens, such as planktonic forms of *C. albicans* [58] and *P. falciparum* (IC_50_ 0.07 µM) [20]. This strong rationale, added to the favorable pharmacological (IC_50_ 2.6/2.8 µM; SI > 35/37) and physicochemical (Fig 2) profiles of MMV675968, motivated the SAR study of its analogs.

The antitrypanosomal pharmacomodulation study of MMV675968 (**Table 4**) revealed that activity varied consistently according to the various substituents added to the core 2,4-diaminoquinazoline structure and the adjacent phenyl portion. For instance, a change in the position of the methoxy substituents in analogs **2, 3, 4** and **5** did not significantly influence (P˃0.05) the activity compared to parent hit **1**. Chlorine removal from the core structure appeared to be beneficial for antitrypanosomal activity and selectivity, as the resultant nonchlorinated analogs **6, 7, 8,** and **9** were more potent and selective than their corresponding chlorinated congeners **2, 3, 4,** and **5**. Of note, this chlorine withdrawal on the core structure led to the identification of analog **7** (6-(((2-methoxyphenyl)amino)methyl)quinazoline-2,4-diamine), which was ∼43-fold and ∼11-fold more active and selective, respectively, than the parent hit **1**. Additionally, analog **7** was ∼53-fold more active and ∼64-fold more selective than the closest analog **3.** Conversely, chlorination of the phenyl portion of the parent hit (**1**) at positions 2 and 5 led to analog **10** (6-(((2,5-dichlorophenyl)amino)methyl)quinazoline-2,4-diamine) that exhibited outstanding ∼58-fold and ∼46-fold increases in activity (IC_50_ 45 nM) and selectivity (SI ∼1737) P˂0.0001). Previously, **Iwatsuki et al** [59] reported an improved antitrypanosomal activity by up to 54-fold for chlorinated antibiotic derivatives compared to nonchlorinated analogs. However, it is noteworthy that depending on the chlorination point on the core or adjacent portion, the chlorine atoms may confer antitrypanosomal activity to the afforded compound. In this line, compound 7 is the most attractive in terms of lipophilic efficiency. Indeed, swapping one methoxy for 2 chlorine substituents in compound 10 increase lipophilicity and might explain the marginal improvement in potency. On another note, polarizing the bond between the phenyl ring and the quinazoline core resulted in a drastic loss in activity (P˂0.0001), as evidenced by increased IC_50_ and mild selectivity values for analogs **11-15** (IC_50_ 2.5 to ˃10 µM) when compared to the profile of close analogs **2, 3, 4** and **5**. Finally, complete loss of activity (IC_50_ ˃10 µM) was observed following shrinkage of the quinazoline core in favor of pyrimidine (Analogs **16-24**). The discrepancies observed in the pharmacological properties of the 23 analogs of MMV675968 indicate that any modification around the quinazoline core may redefine its binding properties to the parasitic target of interest. Analogs **7** and **10** emerged from the SAR study as the more promising candidates. They were therefore prioritized and progressed for additional analyses.

Further studies on analogs **7** and **10** included deciphering their time kill kinetics, induction of intracellular ROS production, membrane permeabilization, DNA fragmentation and ferric ion reducing antioxidant power (FRAP). Compound **7** was found to alter the growth of parasites after a period of 24 hours at IC_99_ and presented a parasitostatic effect on trypanosomes, which was further confirmed by the reversibility of the drug effect after drug removal and subculture. We can therefore argue that this compound is either a slow-acting inhibitor or might be involved in a reversible interaction with the trypanosome metabolic target of action. Such a drug might rapidly induce drug resistance selection and will require either multiple and high doses to cure the mice in an *in vivo* study or a very long treatment period. This profile of compound **7** is of limited advantage compared to the current remedies used for the treatment of trypanosomiasis. Of particular interest, compound **10** presented a fast-killing effect within 4h and irreversible cidal activity during the whole monitoring period of 72h at its IC_99_, contrary to compound **7,** and pentamidine (reference anti-trypanosomiasis drug) exhibited a gradually diminishing effect from 24-72h postdrug removal. This time-kill kinetics profile of compound **10** validates it as a promising candidate for further development against trypanosomiasis.

Further attempts to understand the mode of action of compounds **7** and **10** against trypanosomes demonstrated that none of the compounds elicited deterrent effects on the plasma membrane permeability of the parasite. Additionally, no induction of a significant imbalance in intracellular ROS levels was observed compared to untreated parasites. Therefore, we suggest that membrane permeability and oxidative stress do not contribute to the mechanism of action of compounds **7** and **10** against *Trypanosoma brucei brucei* parasites.

Moreover, exploration of the mode of elicited cell death indicated that the compounds did not induce cell death through cytolysis, as no DNA fragments were detected in the cell supernatant. This finding corroborates the absence of a deterrent effect by compounds **7** and **10,** as previously demonstrated in our membrane permeability assay. Subsequent detection of DNA fragments in the cell lysate of parasites exposed to compound **10** confirmed its apoptosis-inducing effect through elicitation of DNA fragmentation in treated bloodstream trypanosomes materialized by a significant increase in absorbance compared to the negative control. Similarly, the positive control (pentamidine) displayed a high DNA fragment signal. Of note, pentamidine has been previously reported to have specific and strong DNA-binding properties, particularly to the minor groove of AT-rich regions [60]. This finding further validates the approach using the DNA fragmentation ELISA kit in our study. Moreover, many other reports have previously mentioned the apoptosis induction potency of pentamidine against Kinetoplastidae [61, 62]. Knowing that DNA fragmentation is the last event of programmed cell death, we can conclude that the type of cell death triggered by compound **10** involves apoptosis. Further investigations are warranted to obtain more insights into the apoptosis induction pathway in trypanosomes exposed to compound **10**. More specifically, it is important to determine whether the inhibitor acts on the kinetoplast or on nuclear trypanosomal DNA and to determine the different biochemical and morphological changes that occur in parasites treated with compound **10**.

Finally, the assessment of the ferric iron reducing ability of the inhibitors showed moderate reducing power by compound **7,** while compound **10** showed no activity. Iron is a vital element in most living organisms, including trypanosomes, and is involved in several important biological processes, such as mitochondrial respiration, DNA replication, antioxidant defense, and glycolysis. In fact, three enzymes were described as being iron-dependent and indispensable for trypanosomes. This is the case for superoxide dismutase, which eliminates superoxide radicals released during generation of the tyrosyl radical in the R2 subunit of ribonucleotide reductase [63, 64]. Alternative oxidase is an important enzyme for the reoxidation of nicotinamide adenine dinucleotide (NADH) produced during glycolysis [65, 66]. In addition, **Ayayi et al.** [67] investigated the iron dependence of oxidase alternative and terminal trypanosomes (AOT) by chelating iron using o-phenanthroline, which resulted in strong inhibition of this enzyme. Ribonucleotide reductase is another iron-dependent enzyme that catalyzes the reduction of ribonucleotides to deoxyribonucleotides needed for DNA synthesis [68, 69]. Therefore, iron deprivation of parasites by compound **7** might induce a loss of viability of these vital enzymes, thereby resulting in a rapid decrease in DNA synthesis, increased oxidative stress and cessation of electron transfer to the AOT enzyme, thus contributing to the death of the parasite.

## Materials and Methods

### Compounds handling and storage

The MMVPB manufactured by Evotec (USA) was obtained free of charge from the MMV (Geneva, Switzerland). The box consisted of 400 drug-like compounds, shipped on dry ice and supplied as five 96-well microtiter plates containing 10 µL of 10 mM stock solutions of compounds in 100% dimethyl sulfoxide (DMSO). Supporting information for compounds was found at https://www.mmv.org/mmv-open/pathogen-box and included plate layout, chemical structures and formula, molecular weights, *in vitro* and *in vivo* DMPK, confirmed biological activities against some neglected disease pathogens and cytotoxicity data. Compounds were diluted to five subsets for a final intermediary concentration of 100 µM in 96-well storage plates using incomplete IMDM (Iscove’s modified Dulbecco’s medium) culture medium (2 µL of stock solution added to 198 µL of sterile incomplete medium). Plates were stored at -20^°^C until biological assays.

The diaminoquinazoline solid analogs were also provided by the MMV organization (Geneva, Switzerland) and were dissolved in DMSO to reach a concentration of 10 mM.

Pentamidine isethionate (Sigma Aldrich) and podophyllotoxin (Sigma Aldrich) were weighed and dissolved in 100% DMSO to a final concentration of 10 mM and further diluted and used as positive controls for the antitrypanosomal and cytotoxicity assays, respectively.

### Antitrypanosomal screening of the Open Access MMV Pathogen Box

#### Parasite growth conditions

The parasite used for this study was the bloodstream form trypomastigotes of *Trypanosoma brucei* subsp*. brucei*, Strain Lister 427 VSG 221 kindly donated by BEI resources (https://www.beiresources.org/). Parasites were axenically cultivated in sterile vented flasks containing complete Hirumi’s modified Iscove’s medium 9 (HMI-9) [500 mL IMDM (Iscove’s modified Dulbecco’s medium) (Gibco, USA) supplemented with 10% (v/v) heat inactivated fetal bovine serum (HIFBS) (Sigma Aldrich), 10% (v/v) serum plus (Sigma Aldrich), HMI-9 supplement (1 mM hypoxanthine, 0.16 mM thymidine, 50 µM bathocuproine disulfonic acid, 1.5 mM cysteine, 1.25 Mm pyruvic acid, 0.2 mM 2-mercaptoethanol (Sigma Aldrich)), and 1% (v/v) penicillin–streptomycin (Sigma Aldrich) and incubated at 37°C in a 5% CO_2_ atmosphere. Cultures were routinely monitored every 72 h using a Lumascope LS520 inverted fluorescence microscope (Etaluma, Inc., USA) to assess parasite density and subsequently passaged with fresh complete medium in such a way that the cell density never exceeded 2 x 10^6^ cells.mL^-1^. [70].

#### *In vitro* single point and concentration–response antitrypanosomal screening

The *in vitro* inhibitory potency of the 400 MMVPB compounds against bloodstream forms of *Trypanosoma brucei brucei* was evaluated using the resazurin-based inhibition assay as previously described [71]. Briefly, parasites at their mid-logarithmic growth phase were counted, and the cell density was adjusted with fresh complete HMI-9 medium to 2 x 10^5^ trypanosomes per mL. Ninety microliters of parasite suspension were then distributed into the wells of 96-well flat-bottomed plates containing 10 µL of compounds for a final test concentration of 10 µM. The first and last columns in each plate served as negative (cells with 0.1% DMSO) and positive (cells with 10 µM pentamidine isethionate) controls, respectively. After 68 hours of incubation at 37°C and 5% CO_2_, parasite viability was checked after fluorescence measurement using a Tecan Infinite M200 fluorescence multiwell plate reader (Austria) at wavelengths of 530 nm for excitation and 590 nm for emission following a 4-hour incubation period with resazurin (0.15 mg/mL in DPBS, Sigma–Aldrich) in darkness. Each assay plate was set up in duplicate and repeated two times. The percent parasite inhibition was determined for each compound based on fluorescence readouts relative to the mean fluorescence of negative control wells.

Compounds exerting a mean inhibition percentage greater than 90% at 10 µM were selected and tested in duplicate at 5-point concentrations using the aforementioned conditions. Likewise, analogs of a selected hit were also tested. Mean fluorescence counts were normalized to percent control activity using Microsoft Excel, and the 50% inhibitory concentrations (IC_50_) were calculated using Prism 8.0 software (GraphPad) with data fitted by nonlinear regression to the variable slope sigmoidal concentration–response formula

y = 100/[1+ 10^(logIC50/99-*x*)*H*^], where *H* is the hill coefficient or slope factor (Singh and Rosenthal, 2001). Prioritized compounds (IC_50_ < 4 µM) were further tested for their cell cytotoxic effect as described below.

### Determination of the cytotoxicity of inhibitors against Vero cells

#### Maintenance of mammalian cells

The African green monkey kidney Vero cell line (ATCC CRL-1586) was grown in T-25 vented cap culture flasks using complete Dulbecco’s modified Eagle’s medium (DMEM) supplemented with 10% FBS, 1% nonessential amino acids and 1% (v/v) penicillin– streptomycin and incubated at 37°C in an atmosphere containing 5% CO_2_. The medium was renewed every 72 h, and cell growth was assessed using an inverted microscope (Lumascope LS520). Subculture was performed when the cells reached ∼80-90% confluence by detaching with 0.25% trypsin-EDTA followed by centrifugation at 1800 rpm for 5 min. The resulting pellet was resuspended and counted in a Neubauer chamber in the presence of trypan blue to exclude nonviable cells colored blue. Once the cell load was estimated, they were either used for the next passage in a new flask or processed for the cytotoxicity assay.

#### Assessment of the cytotoxic effect of compounds

The cytotoxicity of promising compounds was assessed as previously described by **Bowling et al.** [71] in a 96-well tissue culture-treated plate. Briefly, Vero cells at a density of 10^4^ cells per well were plated in 100 µL of complete DMEM and incubated overnight to allow cell attachment. Plates were then controlled under an inverted fluorescence microscope (Lumascope LS520) to assure adherence, sterility and cell integrity. Thereafter, culture medium from each well was carefully emptied, and plates filled with 90 µL of fresh complete medium followed by the addition of 10 µL of serial 5-fold dilutions of compound solutions. Podophyllotoxin (100 µM-0.16 µM) and 0.5% DMSO (100% cell viability) were also included in assay plates as positive and negative controls, respectively. After an incubation period of 48 h at 37°C in a humidified atmosphere and 5% CO_2_, 10 µL of a stock solution of resazurin (0.15 mg/mL in DPBS) was added to each well and incubated for an additional 4 h. Fluorescence was then read using a Magelan Infinite M200 fluorescence multiwell plate reader (Tecan) with excitation and emission wavelengths of 530 and 590 nm, respectively. The percentage of cell viability was calculated from readouts, and the median cytotoxic concentration (CC_50_) for each compound was deduced from concentration–response curves using GraphPad Prism 8.0 software as described above. Selectivity indexes were then determined for each test substance as follows:

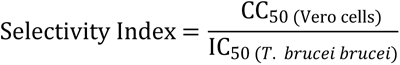

### *In silico* exploration of the DHFR and TR binding properties of analogs 7 and 10

#### Compound preparation

Compound structures were drawn and converted to SMILES format using the smiles generator window of the cheminfo webserver (http://www.cheminfo.org/). Accessible protonation states of the compounds at physiological pH were assessed using the dimorphite-DL program [33]. Dimorphite-DL uses a straightforward empirical algorithm that leverages substructure searching and makes use of a database of experimentally characterised ionizable molecules to enumerate small-molecule ionization states [33]. Here the SMILES format of the compounds was used as input, generating a list of SMILES of all the possible protonation states at the default physiological pH (6.4 – 8.4). The output from dimorphite-DL was streamlined by measuring the pKa values using MolGpka [34]. The MolGpka webserver predicts pKa through a graph-convolutional neural network model that works by learning pKa related chemical patterns automatically and building reliable predictors with the learned features [34]. Matching the pKa values of the different ionizable sites with the physiological pH resulted in the retention of only the physiologically feasible compounds. The retained SMILES were then converted to the vina compatible pdbqt format using the format converter window of the cheminfo webserver in preparation for docking.

#### Protein structure preparation

Protein structures used in this study were obtained from the Research Collaboratory for Structural Bioinformatics, Protein Data Bank (RCSB PDB) [72]. For *Trypanosoma brucei brucei* trypanothione reductase, the crystal structure (PDB ID: 2WP6) is available and was downloaded. On the other hand, *T brucei brucei* DHFR crystal structure has not been deposited in the RCSB PDB. However, it shares 100% sequence identity with *T brucei rhodesiense* DHFR, hence the latter’s crystal structure (PDB ID: 3RG9) was downloaded and prepared for use. The protein structures were pre-processed using Discovery studio visualizer version 4.1 [73]. Initially the structure of DHFR was examined for the presence of the recently identified DHFR crystal structural error, reported in *P falciparum* [74] . The implicated loops were identified to be shorter and capable of no entanglements, hence ruling out the possibility of a crystallographic error. The cofactors NADPH and FAD were maintained in both the DHFR and TR structures respectively. Partial charges and AutoDock atom-types (pdbqt format) were incorporated in the protein and cofactor structures respectively using the *prepare_receptor4.py* and the *prepare_ligand4.py* python scripts from AutoDock4 tools [75].

#### Molecular Docking

A blind docking protocol was implemented using the molecular docking and virtual screening tool AutoDock Vina [76]. Docking validation was accomplished through a redocking of the co-crystalized ligands of the DHFR and TR structures into their respective active sites. Docking parameters adopted following the validation were as follows: DHFR - box size (in Å) x = 42.75, y = 42.75, z = 41.62; box centre x = 64.64, y = 32.86, z = 36.54; and an exhaustiveness of 290. TR - box size (in Å) x = 83.62, y = 65.62, z = 84.00; centre x = 43.17, y = 5.75, z = - 0.05; and an exhaustiveness of 290. After the docking a split of the top nine predicted poses from AutoDock vina was performed using the *vina_split* script and the pose with the lowest docking score was retained for further evaluations. Protein – ligand complexes were then prepared for the top scoring ligands using an inhouse python script and visualisation was done in Discovery studio visualizer version 4.1 and PyMOL [77].

#### Molecular dynamics simulations

Molecular dynamics (MD) simulations were performed within the Amber forcefield a99SB-disp [78], using the GROMACS v.2018 software package [79]. The GROMACS compatible version of the a99SB-disp forcefield was obtained from [80] and used to perform all-atom MD simulations. Ligand parameters were determined by the ACPYPE tool [81]. TIP4P (a99SBdisp_water) water molecules were used to embed each system within a cubic simulation box, leaving a clearance space of 1.0 Å from the edges of the protein. Appropriate amounts of Na+ and Cl-ions were added to neutralise the total system charges. This was followed by system relaxation through energy minimisation using the steepest descent algorithm with a force threshold of 1000 kJ/mol/nm and a maximum of 50,000 steps. The temperature and pressure were respectively equilibrated using the modified Berendsen thermostat (at 300 K for 100 ps), according to the NVT ensemble and the Parrinello–Rahman barostat [82], according to the NPT ensemble, to maintain the pressure at 1 bar. During the equilibration steps, the protein was position restrained and constraints were applied to all the bonds using the LINCS algorithm [83]. Finally, unrestrained production runs were performed for 100 ns each under periodic boundary conditions (PBC) and the equilibration step thermostat and barostat were both maintained for temperature and pressure couplings. The leap-frog integrator was used with an integrator time step of 2 fs, while the Verlet cut-off scheme was implemented using default settings, and coordinates were written at 10.0 ps intervals. Long-range electrostatic interactions were treated using the Particle-mesh Ewald (PME) algorithm [84], while short-range non-bonded contacts (Coulomb and van der Waals interactions) were defined at a 1.4 nm cut-off. All the analyses were accomplished using GROMACS tools. The trajectories were first corrected for periodic boundary conditions using the *gmx trjconv* tool - starting with system centring within the simulation box, fitting the structures to the reference frame, and putting back atoms within the box. Furthermore, *gmx rms*, *gmx rmsf*, and *gmx gyrate* were utilized for the calculation of the root mean square deviation (RMSD), root mean square fluctuation (RMSF), and radius of gyration (Rg) respectively. The *gmx_cluster* tool was used for ligand clustering, and the gromos method was used with an rmsd cut-off value of 0.12 nm.

### Attempts to elucidate other antitrypanosomal modes of action of analogs 7 and 10

#### *In vitro* determination of parasite-killing kinetics of selected hit compounds

The preliminary structure-activity-relationship study of compound MMV675968 (**1**) led to the identification of two highly potent and selective hits (analogs **7** and **10**). Thus, they were selected, and their effect at various concentrations was assessed on the proliferation rate of the bloodstream form of *Trypanosoma brucei brucei* using microscopic cell count. Briefly, parasites at their exponential growth phase were seeded into a 24-well flat-bottomed plate and incubated for 72 h at a cell density of 2 x 10^5^ trypanosomes per mL with compounds at their IC_99_, IC_90_, IC_50_, and IC_10_ values. After specified exposure time intervals (0, 4, 8, 12, 24, 30, 36, 48, 60, 72 h), the content of each well was harvested and counted on a Neubauer hemacytometer to determine the number of mobile parasites [85]. The obtained values were used to plot the growth curves as parasite density (cells/mL) versus incubation time using GraphPad Prism 8.0 software. All experiments were performed in duplicate and included positive (pentamidine) and negative (parasites without inhibitor) controls.

#### Assessment of the cidal or static effect of the inhibitors

Data generated from the time-kill kinetics indicated that analogs **10** and **7** at their respective IC_99_ totally inhibited the growth of trypanosomes within 4 and 24 hours, respectively. Therefore, the ability of parasites to recover post exposure to inhibitors **10** and **7** was further evaluated. Briefly, parasites at 2 x 10^5^ trypanosomes/mL were incubated at 37°C and 5% CO_2_ with compounds at their respective IC_99_ and IC_90_ concentrations for 4 hours and 24 hours for compounds **10** and **7,** respectively. After that, the cells were washed three times with fresh complete medium by centrifugation at 2500 rpm for 7 minutes. Cell pellets were thereafter resuspended in fresh complete culture medium in a 24-well plate and subcultured under the same conditions for 68 hours for compound **10** and 48 hours for compound 7 to achieve the complete parasite cycle. Thereafter, cells were enumerated using the Neubauer cell counter at each time point. The assay was performed in duplicate, and the mean cell counts were plotted membrane against time using GraphPad Prism 8.0 to assess the cidal or static effect of inhibitors.

#### Effect of compounds 7 and 10 on trypanosome plasma membrane integrity

The plasma membrane is the first barrier protecting the cell from the external environment and therefore might represent an important challenge for compounds to exert their inhibitory action. Thus, compounds **7** and **10** were assessed for their ability to induce alterations in cell integrity as previously described [85] with a modification consisting of using a SYBR green assay [86]. The principle of this assay is based on the fact that upon treatment with membrane disruptors, trypanosomes with compromised plasma become fluorescent as a result of SYBR green entry and fixation to the exposed DNA. Practically, in a 96-well microtiter plate, 90 µL of trypanosome suspension (2 x 10^6^ cells/mL/well) was preincubated with SYBR Green (2X) (Sigma Aldrich) for 15 min at 37°C and 5% CO_2_ in darkness. The reaction was then allowed to start following the addition of 10 µL of compounds at their IC_99_, IC_90_, IC_50_ and IC_10_. The plate was then incubated at 37°C on a Magelan Infinite M200 multiwell plate reader (Tecan), and fluorescence was further recorded every 5 min for up to 120 min at λex = 485 and λem =538 nm. Wells containing saponin (Sigma Aldrich) at 0.075 g/mL as a positive control for maximal permeabilization, untreated trypanosomes and HMI-9 medium representing the negative control and the background signal, respectively, were also included. The results are expressed as the mean±SD of two experiments carried out in duplicate after deducting the background signal (wells containing HMI-9 medium) and then used to plot the fluorescence counts versus time (min) graph.

#### Induction of intracellular reactive oxygen species (ROS) production by trypanosomes upon treatment with inhibitors

The production of ROS was detected using a 2’,7’–dichlorofluorescein diacetate (DCFDA) probe (Sigma–Aldrich) as described by **Rea et al.** [87]. Briefly, parasites (2 x 10^6^ cells/mL/well) were washed in incomplete IMDM medium and incubated with compounds **7** and **10** at their respective IC_99_, IC_90_, IC_50_ and IC_10_ for 2 hours at 37°C. DCFDA (100 µL, 5 µM) was then added to the parasites, and incubation was pursued for 15 min at 37°C. Hydrogen peroxide (H_2_O_2_ 0.1% v/v) was used as a positive control for maximal ROS production, and wells with untreated parasites were included as a negative control. Of note, DCFDA is cleaved by ROS to produce fluorescent 2′,7′-dichlorofluorescein (DCF), of which the fluorescence intensity was measured at λex = 485 nm and λem = 520 nm. Data obtained from duplicate readouts were used to determine the ROS production percentage relative to the positive control (100% Production).

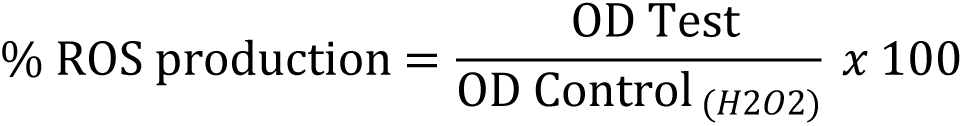

#### Induction of cellular DNA fragmentation upon trypanosome treatment with inhibitors

In an attempt to understand the tentative mechanism (apoptosis or cytolysis) of trypanosome death following exposure to compounds **7** and **10**, DNA fragmentation analysis was performed using a cellular DNA fragmentation ELISA kit according to the manufacturer’s recommendations **(ROCHE Cat. No. 11585045 001 Sigma Aldrich).** Basically, this assay tends to quantify apoptotic cell death by detection of fluorescent dye-labeled DNA fragments in the cytoplasm of affected cells or to measure cell-mediated cytotoxicity by detection of dye-labeled DNA fragments released from damaged cells into the culture supernatant. Briefly, parasites (4 x 10^5^ parasites/mL/well) were incubated at 37°C for 24 hours with BrdU (5’-bromo-2’-deoxyuridine), a nonradioactive thymidine analog that is incorporated into genomic DNA. BrdU-labeled parasites were harvested (by spinning at 2500 rpm for 7 min) followed by centrifugal washing using BrdU-free HMI-9 medium. One hundred microliters of the labeled and washed parasites at a final density of 4 x 10^5^ parasites/mL/well were treated in duplicate wells of a 96-well round-bottom plate with 100 µL of analogs **7** and **10** at their respective IC_99_, IC_90_, IC_50_ and IC_10_ for 4 hours. The plate was then centrifuged at 2500 rpm for 7 min, and the pellets were collected and kept at 4°C until further use. For cytolysis measurement, the supernatants were added to a microplate coated with anti-DNA antibody to allow DNA capture from the test samples. The captured DNA fragments were subsequently denatured using microwave irradiation (LG NeoChef Charcoal Healthy Ovens) at 500 W for 5 minutes to separate DNA strands in view of displaying the BrdU. Thereafter, an anti-BrdU-antibody-POD (peroxidase) conjugate was added to detect the BrdU contained in the captured DNA fragments. One hundred microliters of the POD substrate solution was added to each well, and the absorbance was read at 370 nm every 30 s until color development, which was green in the case of this study.

For apoptosis measurement, the cell pellets from each well were resuspended in 200 µL of the kit’s incubation buffer and incubated for 30 min. The lysed cells were centrifuged at 1700 rpm for 10 min, and the supernatants were used as described above. Triton X-100 (Sigma Aldrich) was used as a positive control for both apoptosis and cell-mediated cytotoxicity. The absorbance values were proportionally correlated to the amount of DNA fragments in the treated cultures. Data are expressed as the mean±SD of two independent experiments performed in duplicate and compared to the negative control (untreated parasites) at a significance level of p˂0.05.

#### Determination of the Ferric Ion Reducing Antioxidant Power (FRAP) of Inhibitors

Iron is one of the vital elements for all parasites, including trypanosomes, as it plays a key role in pathogenesis and the host immune response. Therefore, iron bioavailability reducing agents to the parasite could be potential candidates for drug development against trypanosomiasis. To this end, we determined the capacity of the test compounds to reduce iron from ferric to ferrous status using the method described by **Benzie et al.** [88]. Briefly, 25 µL of each compound (**7** and **10**) was added to 25 µL of a solution of Fe^3+^ prepared at 1.2 mg/mL in distilled water in a 96-well microplate. The plate was incubated for 15 min at room temperature in darkness, after which 50 µL of ortho-phenanthroline (0.2% in methanol) was added to achieve final compound concentrations ranging from 400 µM to 0.19 µM. Plates were reincubated for 15 min, and the absorbance was measured at 510 nm. Ascorbic acid was used as a positive control and tested at concentrations ranging from 100 to 0.048 μg/mL. The median reducing concentration (RC_50_) values of compounds were determined through sigmoidal concentration–response curves using GraphPad Prism version 8.0 software.

#### Statistical analysis

Data collected from at least two independent experiments performed in duplicate are expressed as the mean ± SD (standard deviation). They were analyzed using Tukey’s multiple comparison test using GraphPad 8.0 software. Differences were considered statistically significant at P˂0,05 (*), P˂0.001 (**), and P˂0.0001 (***).

## Conclusion

Two promising 2,4-diaminoquinazoline analogs [**MMV1578445 (10)** and **MMV1578467 (7)]** with potent antitrypanosomal activity and high selectivity toward Vero cells emerged from our study. In light of the above evidence, their mechanism of antitrypanosomal action does not include membrane permeabilization or oxidative stress generation. Nevertheless, compound **MMV1578445 (10)** was found to induce *Trypanosoma* death through DNA fragmentation as a sign of late apoptosis, while compound **MMV1578467 (7)** showed a moderate ferric iron reducing ability. Both compounds are potent binders of DHFR, eliciting important diaminoquinazoline ring mediated interactions with key DHFR ligand recognition residues including Val32, Asp54, and Ile160. Owing to their favorable pharmacological and physicochemical properties, compounds **7** and **10** qualify as suitable starting points for the development of alternative treatments for trypanosomiasis. Hence, we plan to further explore their pharmacological properties and mechanisms of action toward validation as drug candidates.

## Acknowledgment

The authors acknowledge MMV for support, designing and supplying the Pathogen Box and the analogs of MMV675968.

## Supporting information

**S1 Table. Chemical structures of the 25 antitrypanosomal MMVPB hit compounds with known anti-kinetoplastid activity.**

**S2 Table. Chemical structures of the 38 antitrypanosomal MMVPB hit compounds with known potency against other diseases.**

